# Modeling of human *MLH1* and *MLH3* variants in mice reveals novel functions of MutLγ in meiotic DNA repair and the ovarian reserve

**DOI:** 10.64898/2026.10.26.762008

**Authors:** Xinbao Ding, Vertika Singh, John C Schimenti

## Abstract

The mismatch repair proteins MLH1 and MLH3 form the MutLγ complex that is essential for >90% of meiotic crossovers (CO) in mice. Although deletion of either gene causes infertility, we previously found that mouse models of certain human *MLH1/3* missense variants caused elevated aneuploidy and progressive loss of female fecundity despite having a normal ovarian reserve. Here, we report that additional missense and null alleles caused significant pre-pubertal follicle loss independent of alterations to meiotic CO levels. Deletion of the DNA damage checkpoint kinase *Chek2 (Chk2)* partially rescued the oocyte pool of *Mlh1-* and *Mlh3*-deficient mice, indicating that defects in meiotic prophase I triggered oocyte elimination. In spermatocytes, both missense and null alleles caused abnormal γH2AX ‘flares’ on autosomes during late pachynema and diplonema, extending into chromatin loops from the synaptonemal complex. These flares frequently contained unusually large foci of the ssDNA binding protein RPA as well as DNA end processing factors including CtIP, RAD50 and BLM, consistent with abnormal and extensive DNA resection. The results reveal a previously unrecognized meiotic function of MutLγ in preventing DNA hyper-resection of spontaneous DNA breaks or of persistent recombination intermediates in late prophase I. Together with our previous studies, the results show that *MLH1/3* variants can impair fertility through at least three mechanistically distinct routes: severe crossover loss sufficient to activate the metaphase I spindle assembly checkpoint; subthreshold crossover defects that increase aneuploidy and embryo loss; and abnormal late-prophase DNA processing that activates a CHEK2-dependent oocyte quality-control pathway and depletes the ovarian reserve, causing primary ovarian insufficiency (POI). Thus, MutLγ has broader roles in maintaining meiotic genome integrity than can be explained by crossover formation alone.

## Introduction

Primary ovarian insufficiency (POI), a disorder involving premature depletion of the ovarian oocyte reserve, affects approximately 1% of women under 40 years of age (1). Genetic causes account for ∼20-25% of patients (2). Whole exome and whole genome sequencing, coupled with genetic studies, have substantially accelerated the identification of genes contributing to POI, many of which are involved in DNA damage repair, homologous recombination, and meiosis (3). These findings highlight the importance of such genes in establishing the ovarian reserve and maintaining ovarian function, but their potential involvement in the 70% of idiopathic POI cases is unknown.

Meiosis is the cellular event that results in the formation of haploid gametes through two successive cell divisions following a single round of DNA replication. During meiotic prophase I, programmed double-strand breaks (DSBs) initiate homologous recombination repair (HRR) that promotes homologous chromosome pairing. A subset of the DSB repair events yields crossovers (COs) that are crucial for proper disjunction of homolog pairs at the first meiotic division. In mice, the DNA mismatch repair (MMR) genes *Mlh1* and *Mlh3* are required for ∼90% of all “Class I” COs which are subject to interference (4). Consistent with this essential function, null alleles of Mlh*1* and *Mlh3* cause sterility in mice (5, 6). Mutant spermatocytes exhibit premature chromosome separation and arrest in meiosis I, while oocytes progress to metaphase I but show chromosome misalignment, spindle defects, rare progression to anaphase, and failed embryogenesis in those that get fertilized after superovulation (6–8)(9). These severe reproductive phenotypes preclude detailed analysis of more subtle roles for MLH1 and MLH3 in promoting mammalian gamete health and function.

In addition to its role in CO formation, MLH1 is crucial for MMR in somatic cells (10, 11). MMR corrects mismatched nucleotides and insertion/ deletion loops generated during DNA replication, and defective MMR leads to microsatellite instability, mutations, and cancer susceptibility (12). For example, spontaneous mutations in *MLH1* are a major cause of colorectal cancer, and inherited alleles underlie Lynch Syndrome, which predisposes to various cancers, namely colorectal and endometrial (13). In canonical MMR, mismatches are recognized by MutSα (MSH2–MSH6) or MutSβ (MSH2–MSH3), which recruit MutL complexes, including MutLα (MLH1–PMS2), MutLβ (MLH1–PMS1), and MutLγ (MLH1–MLH3), to coordinate downstream repair (10). Thus, MLH1 and MLH3 function at the intersection of meiotic recombination and genome maintenance, though MLH3 is predominantly involved in meiotic recombination; it serves as the catalytic subunit of the MLH1/3 complex that contains an endonuclease function essential for resolution of double Holliday junction intermediates (14, 15).

Due to the importance of *MLH1/3* in cancer prevention, fertility, and genetic recombination, we used mouse modeling to identify pathogenic missense variants in the human population (16). Several of these mouse models - engineered to contain selected missense variants in the corresponding positions of the mouse orthologs - exhibited reproductive defects in one or both sexes. In all 5 analyzed *Mlh1* models, females exhibited markedly decreased litter sizes with age though not all exhibited reduced crossovers or aneuploidy-associated pregnancy loss. Additionally, only one (*Mlh1^K618T^*) exhibited a reduced ovarian reserve around puberty (16), and this allele had about a 50% reduction in COs. Whereas elevated aneuploidy and resulting embryo loss is an expected consequence in mutants with reduced crossovers, the reduced ovarian reserve was surprising. Gamete loss during oocyte meiosis is typically driven by earlier defects in DSB repair or homolog synapsis such as those caused by deficiency of *Spo11* (needed to make DSBs that drive HRR-mediated homolog pairing), *Dmc1* and *Msh5* (HRR genes) (17–19) that subsequently trigger checkpoint-mediated elimination during pachynema (20). Meiocytes deficient for either *Mlh1* or *Mlh3* are not asynaptic and do not have unrepaired meiotic DSBs (5–7), so a decrease in the oocyte pool at a young age is unexpected and likely due to hitherto unrecognized defects that trigger loss prior to dictyate arrest. Regarding the precipitous decline in fecundity with age, we hypothesized that either POI was entirely responsible, or that a substantial fraction of oocytes in the mutants was of poor quality, and that these were selected for maturation only after the pool of higher quality ones became exhausted (16).

To explore the basis for reduced fecundity with age and to determine of other Mlh1/3 variants caused ovarian reserve depletion, we selected and characterized the phenotypes of 4 additional predicted pathogenic missense polymorphisms, three of which caused varying degrees of follicle depletion, even though CO perturbation and aneuploidy was either minor or unaffected in two of them. However, the hypomorphic alleles, along with null mutants in both genes, exhibited persistent unrepaired DNA damage – characterized by large stretches of ssDNA - in the late pachytene and diplotene stages, suggestive of abnormal DNA hyper-resection. Genetic experiments indicate that this damage underlies oocyte elimination through CHK2-mediated activation of the DNA damage checkpoint. These results reveal a previously unrecognized germline protection mechanism mediated by MLH1/3 during late prophase I of meiosis.

## Results

### Generation of knock-in mice bearing human missense variants predicted to be pathogenic

As part of a project to experimentally evaluate potential infertility variants in the human population, we have focused on *MLH1* and *MLH3* for several reasons important for reproductive health. Whereas severe alleles in these genes cause infertility, milder alleles can lead to medically relevant issues including POI and germ cell aneuploidy (16). To identify candidate missense variants for validation, we established a multi-step prioritization pipeline for *MLH1* and *MLH3* missense variants (Fig. S1A). Variants were evaluated using five *in silico* prediction algorithms (Fig. S1B). Candidates were prioritized if they were predicted to be deleterious by multiple algorithms and had a total allele frequency of <0.1%. Applying these criteria yielded 157 candidate variants in MLH1 and 116 candidate variants in MLH3, from a total of 448 and 760 missense variants identified in the respective genes. We further evaluated the predicted effects of these variants on protein stability using the mCSM algorithm (21). Interestingly, ∼90% of the prioritized variants (91% of MLH1 variants and 88% of MLH3 variants) were predicted to decrease protein stability (ΔΔG < 0), suggesting that protein destabilization may be a common mechanism by which missense variants impair MLH1 and MLH3 function (Fig. S1E). Among these destabilizing variants, 7 were previously modeled in mice by our lab (16, 22), demonstrating the utility of this prioritization strategy. In present study, we selected 4 additional missense variants for *in vivo* analysis: *MLH3 Mlh3^P1262S^*, *MLH3^F120S^*, *MLH1^N64S^*and *MLH1^G98S^* encoded by rs372912259, rs28756979, rs63750952 and rs267607725, respectively (Table 1). Notably, although MLH1^N64S^ was predicted to have little effect on protein stability, there was a report that this variant reduces MLH1 protein abundance by ∼40% *in vitro* (23).

**Table 1.** Predicted functional consequences of human missense variants.

| Gene | Variants | AAS | AF (%) | Functional prediction |  |  |  |  |
| --- | --- | --- | --- | --- | --- | --- | --- | --- |
|  |  |  |  | REVEL | PROVEAN | SIFT | Polyphen2 | CADD |
| MLH3 | rs372912259 | P>S | 0.07079 | 0.959 | -4.64 | 0.01 | 0.983 | 26 |
| MLH3 | rs28756979 | F>S | 0.02589 | 0.873 | -5.28 | 0 | 0.982 | 26.3 |
| MLH1 | rs63750952 | N>S | 0.003536 | 0.926 | -4.41 | 0 | 0.996 | 26.4 |
| MLH1 | rs267607725 | G>S | 0.0004 | 0.996 | -5.45 | 0 | 1 | 32 |

Deleterious threshold scores: REVEL, >0.5; PROVEAN: <-2.28; SIFT: <0.05; PolyPhen2: >0.8; CADD: >25.

**Table 2.** Phenotypic summary of MLH1 and MLH3 missense variants in mouse models.

| GENE | CO DEFECT | ANEUPLOIDY | DNA DAMAGE | FOLLICLE LOSS |
| --- | --- | --- | --- | --- |
| MLH3 P. R93G | No | No | No | No |
| MLH3 P. R1230H | Yes | N.A. | N.A. | N.A. |
| MLH3 P. A1394T | No | No | N.A. | N.A. |
| MLH3 P. F120S | No | No | Yes | Yes |
| MLH3 P. P1262S | No | No | Yes | Yes |
| MLH3 P. V384D | No | No | N.A. | N.A. |
| MLH1 P. E268G | Yes | Yes | N.A. | N.A. |
| MLH1 P. K618E | No | No | N.A. | N.A. |
| MLH1 P. K618T | Yes | Yes | Yes | Yes |
| MLH1 P. K618A | Yes | Yes | Yes | Yes |
| MLH1 P. V326A | No | No | N.A. | No |
| MLH1 P. V716M | Yes | No | N.A. | No |
| MLH1 P. N64S | No | No | Yes | Yes |
| MLH1 P. G98S | N.A. | N.A. | N.A. | N.A. |

N.A., not available

Mice bearing the human amino acid substitutions were introduced into the corresponding positions in the mouse orthologs using CRISPR/Cas9 mediated genome editing in embryos (Fig. S2A), then bred to homozygosity for phenotypic evaluations. *Mlh3^P1262S^*, *Mlh3^F120S^*, and *Mlh1^N64S^* homozygotes were viable (these alleles are abbreviated hereafter as *Mlh3^PS^, Mlh3^FS^* and *Mlh1^NS^,* respectively). By contrast, founder mice carrying the p.G98S substitution in *Mlh1* were infertile; males were azoospermic and female founders failed to produce offspring (Fig. S2B and C). The p.Asn64 and p.Gly98 residues are both located in the ATPase domain of human MLH1 (Fig. S2D). Frameshift alleles of *Mlh1* and *Mlh3* were also recovered and expanded as contemporaneous null models (*Mlh1^-/-^* and *Mlh3^-/-^*) for comparison (Fig. S2C). Additionally, we included a mouse line modeling the *Mlh1^K618A^* (*Mlh1^KA^*) polymorphism, which is located within the EXO1-interaction domain of MLH1. This variant hadn’t been phenotypically characterized beyond confirming that homozygous mice were viable and fertile (16). Together, these knock-in and knockout models allowed us to compare the effects of partial and complete disruption of MLH1/MLH3 function on gametogenesis and ovarian reserve maintenance.

To investigate why *Mlh1^G98S^* knock-in founders were infertile, we modeled ATP/Mg²⁺-bound MLH1 structures for WT, p.G98S, p.N64S, and p.G67R (reported to impair MMR and CO (25)) using AlphaFold 3 (24). Pocket analysis (26) identified G67 and G98, but not N64, within the ATP-binding pocket and predicted that G98S produced the greatest structural divergence from WT, including remodeling of the ATPase pocket and displacement of several canonical ATPase residues (Fig. S3A-D). Although the region surrounding G98 showed relatively low AlphaFold confidence, its altered conformation was reproducible across independent models, and complementary structural analyses predicted disruption of local packing and flexibility (Fig. S3E-G). Collectively, these analyses identify G98S as the variant with the strongest predicted impact on MLH1 ATPase-domain structure, without evidence for loss of ATP-Mg²⁺ binding.

### Differences in severity of spermatogenesis and CO defects in humanized variant alleles

Consistent with originally reported phenotypes of *Mlh1/3* mouse mutants (6, 7), the *Mlh1^-/-^* and *Mlh3^-/-^*mice we generated were sterile and azoospermic, with smaller testes that showed complete maturation arrest (no progression past meiotic prophase I; Fig. S4 and S5A-C). In contrast, fertility testing of the male knock-in models *Mlh3^PS/PS^*, *Mlh3^FS/FS^*, *Mlh1^NS/NS^* and *Mlh1^KA/KA^* all sired offspring when mated with wildtype (WT) females (Fig. S4) (16). Whereas the former 3 had testis weight comparable to controls, *Mlh1^KA/KA^*testes were slightly (∼20%) smaller (Fig. S5A). However, the sperm counts of *Mlh3^FS/FS^* and *Mlh1^KA/KA^* males were significantly reduced to ∼1/2 and 1/3 of controls, respectively (Fig. S5B). Histological examination of testes revealed grossly normal seminiferous tubule architecture and well-organized germ cell distribution in all 4 knock-in models, as well as in *Mlh3^PS/-^*, *Mlh3^FS/-^* and *Mlh1^NS^*/^-^ mice (Fig. S5C). However, abnormal metaphase-like spermatocytes – some with lagging chromosomes – were evident in *Mlh1^KA/KA^*tubules (Fig. S5C). To investigate further, TUNEL (terminal deoxynucleotidyl transferase dUTP nick end labeling) staining was performed on homozygous mutant genotypes. *Mlh1^-/-^*and *Mlh3^-/-^* testis exhibited widespread apoptosis (Fig. S5D), consistent with previous studies (6, 7). There were small but significant increases in *Mlh3^PS/PS^*, *Mlh3^FS/FS^*, and *Mlh1^NS/NS^* mice (Fig. S5D). In addition to *Mlh1^KA/KA^* testes having a substantial increase in TUNEL+ germ cells, phospho-histone H3 (pH3) co-labelling of spermatocytes indicated that the apoptotic cells were largely at metaphase I, consistent with the histological observations (Fig. S5E and F), thus raising the possibility of CO aberrations that might be triggering the spindle assembly checkpoint.

To test this, we quantified MLH1 or MLH3 foci in the mutant lines (using anti-MLH3 antibody for *Mlh1* mutants and anti-MLH1 for *Mlh3* mutants), which serve as proxies for Class I CO sites, comprising about 90% of all COs (6, 7). As expected, *Mlh1^-/-^* and *Mlh3^-/-^*spermatocytes had complete or severe loss of MLH1/MLH3 foci on pachytene chromosomes, and consistent with loss of most COs, had only 1-3 bivalents detectable in metaphase I (MI) chromosome spreads. Focus numbers in *Mlh3^PS/PS^*, *Mlh3^FS/FS^*, and *Mlh1^NS/NS^* spermatocytes were no different than WT and heterozygous counterparts (Fig. 1A and B), and these mutant genotypes revealed a normal number of 20 bivalents (Fig. 1C). However, *Mlh1^KA/KA^*spermatocytes had a significant reduction in MLH1 foci (Fig. 1A and B), as well as increase of 1∼2 metaphase I univalents (p=.0001) (Fig. 1C). These results suggest that the cause of apoptosis seen in the humanized alleles - with the possible exception of *Mlh1^KA^*- is probably not entirely SAC activation.

**Figure 1.**
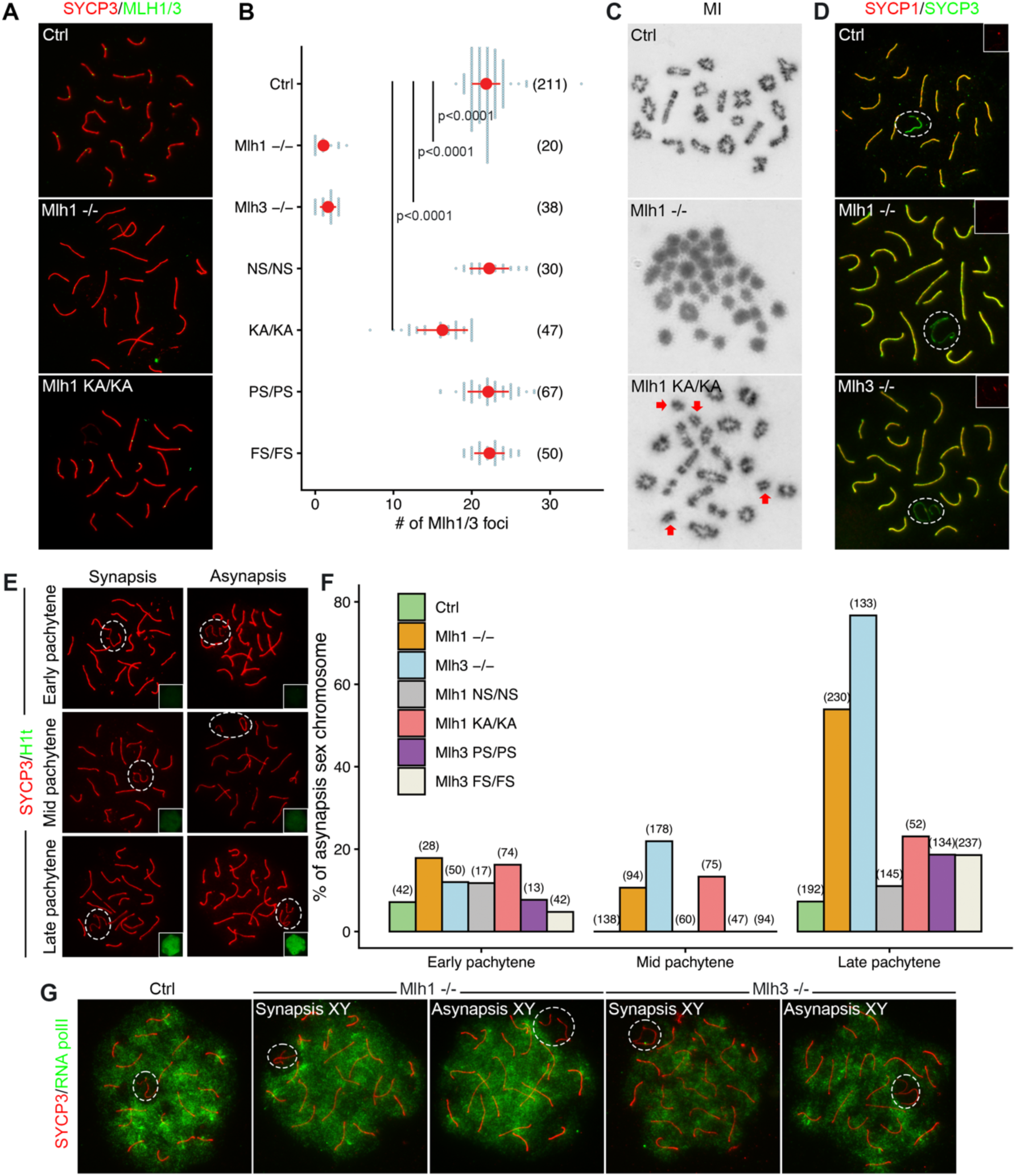
Crossing over and synapsis in *Mlh1* and *Mlh3* mutant spermatocytes. A) Immunostaining of SYCP3 and MLH1 (for *Mlh3^-/-^, Mlh3^PS/PS^*and *Mlh3^FS/FS^* mice) or MLH3 (*Mlh1^-/-^*, *Mlh1^NS/NS^*and *Mlh1^KA/KA^* mice) in surface spread adult spermatocytes. B) Quantification of MLH1 or MLH3 foci. P values for those mutants significantly different from Ctrl (+/+ and heterozygotes) are indicated. Parenthetical numbers = N of spermatocytes, with each represented as light grey dot. C) Meiotic metaphase I spreads from adult Ctrl and mutant spermatocytes. The arrows indicate univalents. D) Immunostaining of SYCP3 and SYCP1 on spermatocyte pachytene chromosome spreads. Insets highlight SYCP1 signals on XY chromosomes. PAR, pseudoautosomal region. E) Representative images of synapsed and asynapsed sex chromosomes at pachytene substages. Insets show nuclear H1t staining, a marker of late pachynema. F) Percentage of asynaptic sex chromosomes at early, mid, and late pachytene. Numbers in parentheses are spermatocytes quantified. G) RNA polymerase II is largely absent from XY bodies. White dashed circles in D, E and G indicate the XY chromosomes. Data in B are represented as the mean ± SEM and were analyzed using one-way ANOVA with Tukey’s *post hoc* test.

To identify other potential defects causing apoptosis, we analyzed chromosome synapsis by co-staining chromosome spreads for SYCP1 and SYCP3. Consistent with previous observations in *Mlh1*^-/-^ spermatocytes (25), autosomal synapsis was largely complete at pachynema, but asynapsis was frequently observed at the pseudo-autosomal region (PAR) of the X and Y chromosomes in *Mlh1*^-/-^ spermatocytes. (Fig. 1D). *Mlh3^-/-^* spermatocytes showed a similar pattern (Fig. 1D). Using histone H1t staining to identify mid-pachytene and later stages of spermatocytes, we quantified the synapsis defects across pachytene substages (Fig. 1E and F). Over 50% of late pachytene *Mlh1^-/-^* and *Mlh3^-/-^* spermatocytes exhibited XY asynapsis, a significantly higher proportion than missense-bearing or control spermatocytes (Fig. 1E and F). Additionally, ∼ 20% of *Mlh3^-/-^*, 10% of *Mlh1^-/-^* and 13% of *Mlh1^KA/KA^* spermatocytes exhibited XY asynapsis at mid-pachynema, which was not observed in the other missense mutants or controls (Fig. 1E and F). These findings suggest that many mutant spermatocytes either lack an obligate Class I CO between the sex chromosomes, thereby preventing or destabilizing synapsis in the PAR (26), or that desynapsis occurs earlier in the mutants. Nevertheless, the defective PAR synapsis in both *Mlh1^-/-^* and *Mlh3^-/-^* spermatocytes did not disrupt meiotic sex chromosome inactivation (MSCI), in that RNA polymerase II was absent from the XY (“sex”) body (Fig. 1G). This is significant in that MSCI disruption is a driver of pachytene arrest (27).

### Follicle loss in *Mlh1/3* mutants

Ovarian gross morphology and histopathology in *Mlh1/3* missense mutants were similar to controls (Fig. 2A), consistent with previous observations of *Mlh1^-/-^* and *Mlh3^-/-^* mice (6, 7). *Mlh3^PS/PS^*, *Mlh3^FS/FS^*, *Mlh1^NS/NS^* and *Mlh1^KA/KA^* females produced offspring (the latter reported previously (16)), whereas *Mlh1^-/-^* and *Mlh3^-/-^* females were infertile (Fig. S4) as expected (6, 7). Despite their apparently normal ovarian histology, the *Mlh3^PS/PS^*, *Mlh1^NS/NS^*, *Mlh1^KA/KA^*, *Mlh1^-/-^* and *Mlh3^-/-^* females had significantly fewer follicles than age-matched controls, particularly at 3 and 8 weeks of age (Fig. 2B and C). The reserve, consisting of primordial follicles, was >50% reduced in *Mlh1^NS/NS^* and *Mlh1^-/-^*, and ∼50% reduced in *Mlh3^-/-^* and *Mlh3^PS/PS^* females. However, the number of growing follicles was not as markedly reduced in the *Mlh3* mutants throughout the range tested (up to 40 weeks), indicating that even with a reduced reserve, similar numbers of primordial follicles were recruited for maturation. Compared to mutants that cause POI via loss of primordial germ cells (PGCs), such as *Mcm9* (28)(29), the depletion was much less severe. Consistent with reduced ovarian reserve, *Mlh3^PS/PS^* and *Mlh1^NS/NS^*females showed age-dependent reductions in litter size over one year of breeding to WT males (Fig. 2D). Overall, the *Mlh3^PS^*, *Mlh1^NS^, Mlh1^KA^* and null alleles of *Mlh1*/*Mlh3* produce a milder follicle loss relative to POI models resulting from PGC defects or meiotic mutants that disrupt chromosome synapsis or DSB repair that completely ablate the oocyte pool (30).

**Figure 2.**
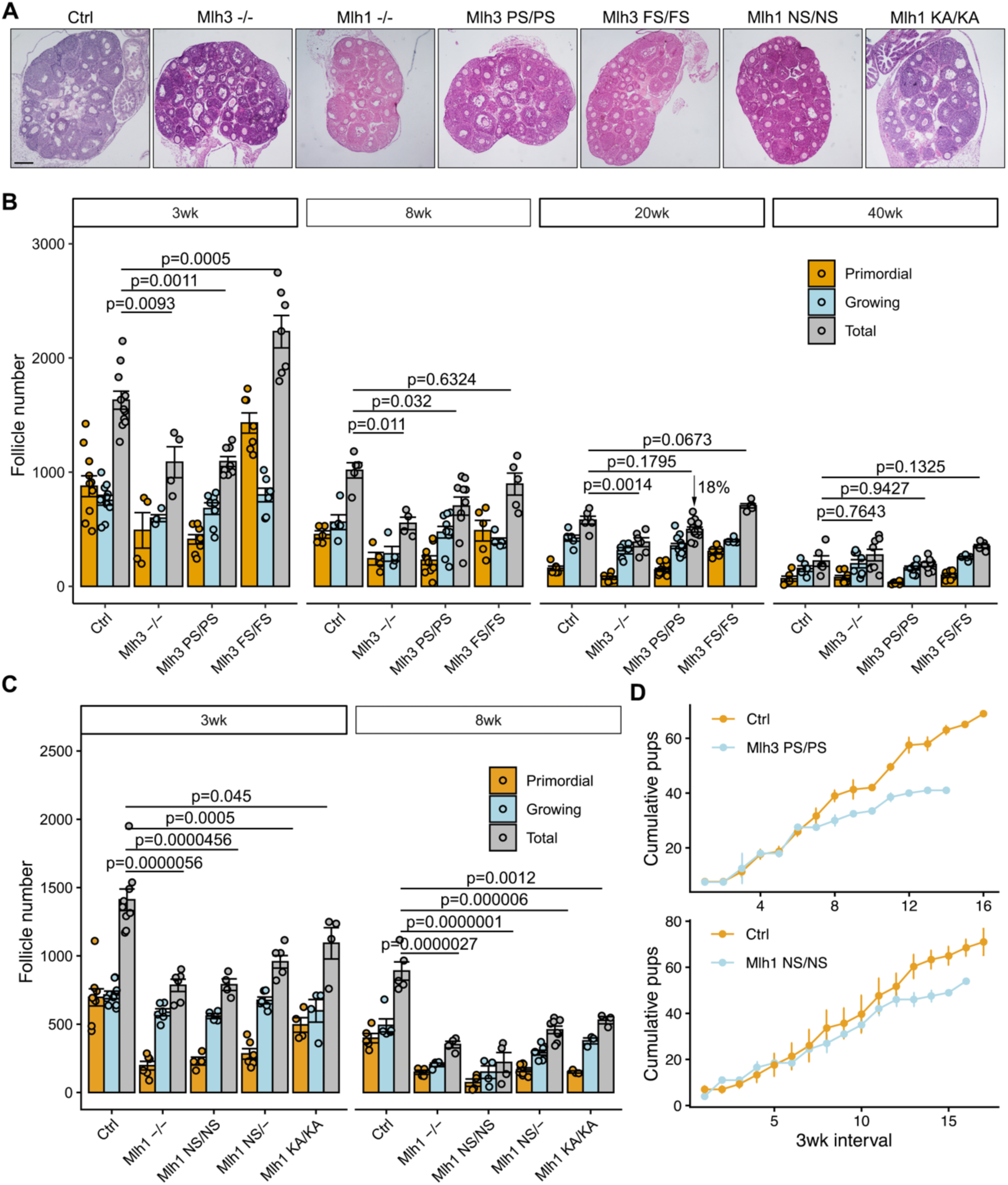
Diminished ovarian reserve in *Mlh1/3* mutants. A) Representative images of 3-wk-old ovaries. Scale bar = 200 μm. B, C) Follicle numbers per ovary from mice of the indicated *Mlh3* (B)and *Mlh1* (C) genotypes and ages (wk = weeks after birth). Each dot represents follicle number in an individual ovary. D) Cumulative number of pups born from fertility trials. The x-axis values are in 3 wk intervals, which approximates litter intervals for fertile females housed continuously with a male. Data in B, C and D are represented as the mean ± SEM, and the total follicle numbers in B and C were analyzed using one-way ANOVA with Tukey’s *post hoc* test.

Because *Mlh1* and *Mlh3* are expressed in somatic and non-meiotic cells, we asked whether null or variant alleles impair early germ cell development. Using an efficient platform for *in vitro* differentiation of embryonic stem cells (ESCs) into primordial germ cell like cells (PGCLCs) (31), we found that multiple *Mlh1* and *Mlh3* missense and null alleles did not significantly affect PGCLC formation (Fig. S6A,B), even though both genes are expressed throughout PGCLC differentiation (Fig. S6C). Additionally, postnatal day 6 *Mlh1^-/-^ and Mlh3^-/-^* testes, in which germ cells have progressed to the spermatogonial phase, showed normal seminiferous tubule morphology without evidence of early germ-cell depletion (Fig. S6D). These results indicate that *Mlh1* and *Mlh3* are dispensable for PGC specification and early postnatal spermatogonial development.

### Persistence of DNA damage in *Mlh1/3* mutant meiocytes

We next sought to determine why *Mlh1/3* variants and nulls reduce the ovarian reserve. Defects in CO number or distribution alone are unlikely to explain the follicle-loss phenotype, as *Mlh3^PS/PS^*, and *Mlh1^NS/NS^*mice have normal numbers of COs (Fig. 1A-C), and we previously reported other variants with declining fecundity and normal CO numbers (16). Additionally, oocyte elimination via the chromosome synapsis checkpoint (32) or MSUC (meiotic silencing of unsynapsed chromosomes) (33–35) seemed unlikely, as no synapsis defects were observed in *Mlh1/3* mutant oocytes (36) or in spermatocyte autosomes (Fig. 1E and F). We then hypothesized that spontaneous DNA damage or meiotic DSBs arising during prophase I might trigger the DNA damage response (DDR) and checkpoint activation, leading to the elimination of oocytes with excessive defects.

The CHK2 kinase (formally, CHEK2) plays a major role in culling mouse oocytes bearing unrepaired meiotic or induced DSBs (37, 38). To test whether *Chk2* is involved in eliminating *Mlh1/3* mutant oocytes, we quantified total follicles in *Mlh3^-/-^ Chk2^-/-^* females compared to single mutants. Loss of *Chk2* significantly increased total follicles in *Mlh3^-/-^*ovaries, reaching a level comparable to WT and single heterozygotes (Fig. 3B). The rescue was due specifically to a dramatic increase in primordial follicles, which constitute the ovarian reserve (Fig. 3B). As expected, *Mlh3^-/-^ Chk2^-/-^*males were azoospermic and their highly achiasmate spermatocytes underwent meiotic arrest, indicative of spindle assembly checkpoint (SAC) activation (Fig. S7). Together, these findings indicate that oocytes contain a form(s) of DNA damage that activates CHK2, causing checkpoint-mediated elimination.

**Figure 3.**
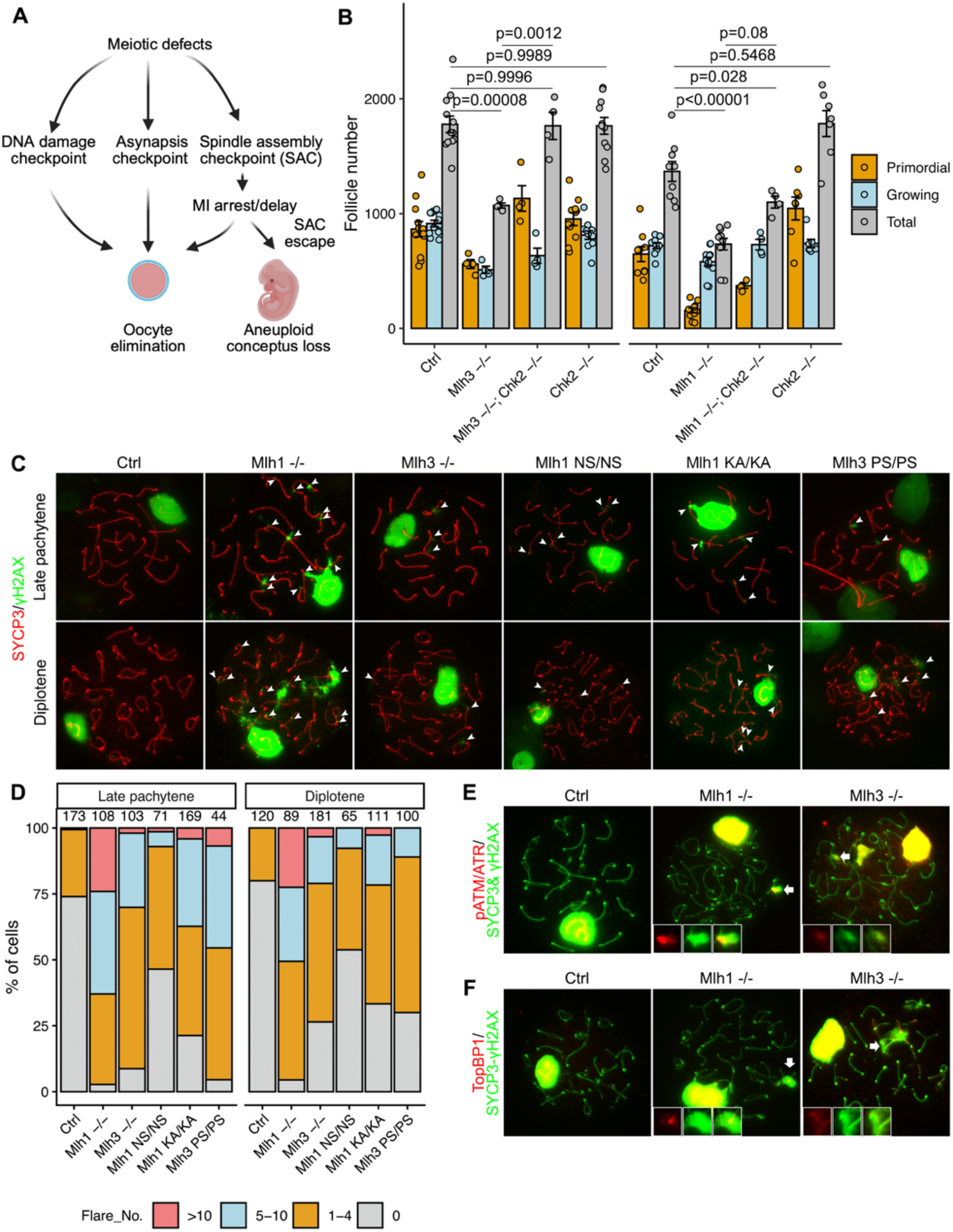
Unresolved DNA damage in late meiotic prophase I *Mlh1/3* mutant spermatocytes. A) Schema illustrating oocyte quality controls. B) Evidence that CHK2 participates in the elimination of *Mlh1* and *Mlh3* deficient oocytes. C) Representative images of γH2AX staining on late pachytene and diplotene chromosome spreads from control (Ctrl; +/+ and heterozygotes) and mutant spermatocytes. D) Quantification and distribution of γH2AX flares on autosomes in mutant spermatocytes. Number above each column indicates the spermatocytes quantified. E) Representative images of ATM/ATR phosphorylated proteins, γH2AX and SYCP3 staining on Ctrl and mutant spermatocytes. F) Representative images of TopBP1, γH2AX and SYCP3 staining of diplotene of Ctrl and mutant spermatocytes. White arrowheads in C indicate the locations of γH2AX flares. The magnified insets in E and F show the γH2AX flares, as indicated by the white arrows. Data in B are represented as the mean ± SEM and were analyzed using one-way ANOVA with Tukey’s *post hoc* test.

We next explored whether *Mlh1/3* mutant meiocytes had evidence of DNA damage in prophase I meiocytes, even though these MMR proteins are not involved in DSB repair and null mutants appear to effectively repair meiotic DSBs (5–7). Spermatocytes were used for this analysis because they are more readily accessible compared to prophase I oocytes, and adult testes contain cells encompassing all stages of meiosis. We immunolabelled spermatocyte chromosome spreads for γH2AX (phosphorylated histone variant H2AX at serine 139), a pan-DNA damage marker. During normal meiotic prophase I, γH2AX is broadly present from leptonema to zygonema and becomes restricted primarily to the sex body as autosomal DSB repair is completed during the pachytene and diplotene stages (39). In *Mlh1/3* mutant spermatocytes, aberrant γH2AX “flares” – projections of staining perpendicular to the SC - were observed on autosomes in late pachytene and diplotene stages (Fig. 3C and D). These were previously observed in late pachytene and diplotene *Mlh1*^-/-^ spermatocytes, but the origins were not explored (25). *Mlh1^-/-^* spermatocytes had the highest frequency of γH2AX flares among the mutants analyzed with >two thirds of cells having 5 or more, whereas most WT cells had none (Fig. 3C and D).

H2AX phosphorylation can occur as a result of transcriptional silencing (as in MSCI or MSUC), DSBs, single-strand breaks/gaps, or ssDNA coated with RPA (replication protein A) (40). Since there was no evidence of autosomal asynapsis, we sought to explore if the flares were associated with various DNA repair pathways. During early meiotic prophase I, DSBs are repaired primarily through homologous recombination, whereas non-homologous end joining (NHEJ) is suppressed. As prophase I progresses, homologous recombination factors decline and NHEJ-associated factors such as 53BP1 begin to appear in late prophase I (21). In both control and mutant spermatocytes, 53BP1 accumulated over sex-chromosome chromatin by mid-pachynema, similar to γH2AX, but in autosomal γH2AX flares in the mutants (Fig. S8A,B), suggesting that these lesions are not marked by canonical NHEJ-associated repair. Late pachytene *Mlh1^-/-^* and *Mlh3^-/-^*spermatocytes showed more persistent axial RPA foci than in controls or knockin mice (Fig. S8C,D). Additionally, control and mutant spermatocytes had similar amounts of RAD51, a RecA homolog which catalyzes strand invasion during DSB repair by HR (Fig. S8E,F).

### *Mlh1/3* mutant spermatocytes accumulate extensive ssDNA within γH2AX flares

Because γH2AX flares were not associated with key NHEJ or HR proteins, we examined other markers of DNA damage intermediates, namely phospho [S/Q]-T sites of ATM/ATR (apical kinases that respond primarily to DSBs and replication errors, respectively) targets, RPA (a single-stranded DNA-binding protein) and TOPBP1 (DNA topoisomerase 2-binding protein 1) which recruits ATR to RPA-coated ssDNA. Both TOPBP1 and ATM/ATR targets were enriched within γH2AX-positive autosomal flares (Fig. 3E and 3F). Normally, RPA broadly localizes along chromosome axes early in prophase I at sites of meiotic DSB resection, then is displaced by recombinases RAD51 and DMC1 as HRR is completed by mid-pachynema. Late-pachytene *Mlh3*^-/-^ and *Mlh1*^-/-^ spermatocytes had elevated RPA foci compared to very low levels in *Mlh3^PS/PS^*, *Mlh1^NS/NS^* and controls (Fig. S8B and C). These RPA foci frequently localized within the γH2AX flares (Fig. 4A). Although overall flare size was similar between mutant and control cells, the RPA-positive regions within flares were significantly enlarged in *Mlh1/3* mutants, resulting in an increased RPA/γH2AX flare size ratio (Fig. 4B). RPA-positive regions within flares were consistently ∼4-fold larger than foci located on the SC (Fig. 4C). Given that the average length of meiotic ssDNA resulting from DSB resection is estimated to be approximately 1,100 nt (41), these measurements suggest that the ssDNA tracts within flares range from ∼200 nt to 12 kb, with an average length of 3.8 kb (Fig. 4C). The average size of RPA-positive flares was greater in diplotene vs. late pachytene spermatocytes, suggesting continued generation of ssDNA during meiotic progression (Fig. 4C).

**Figure 4.**
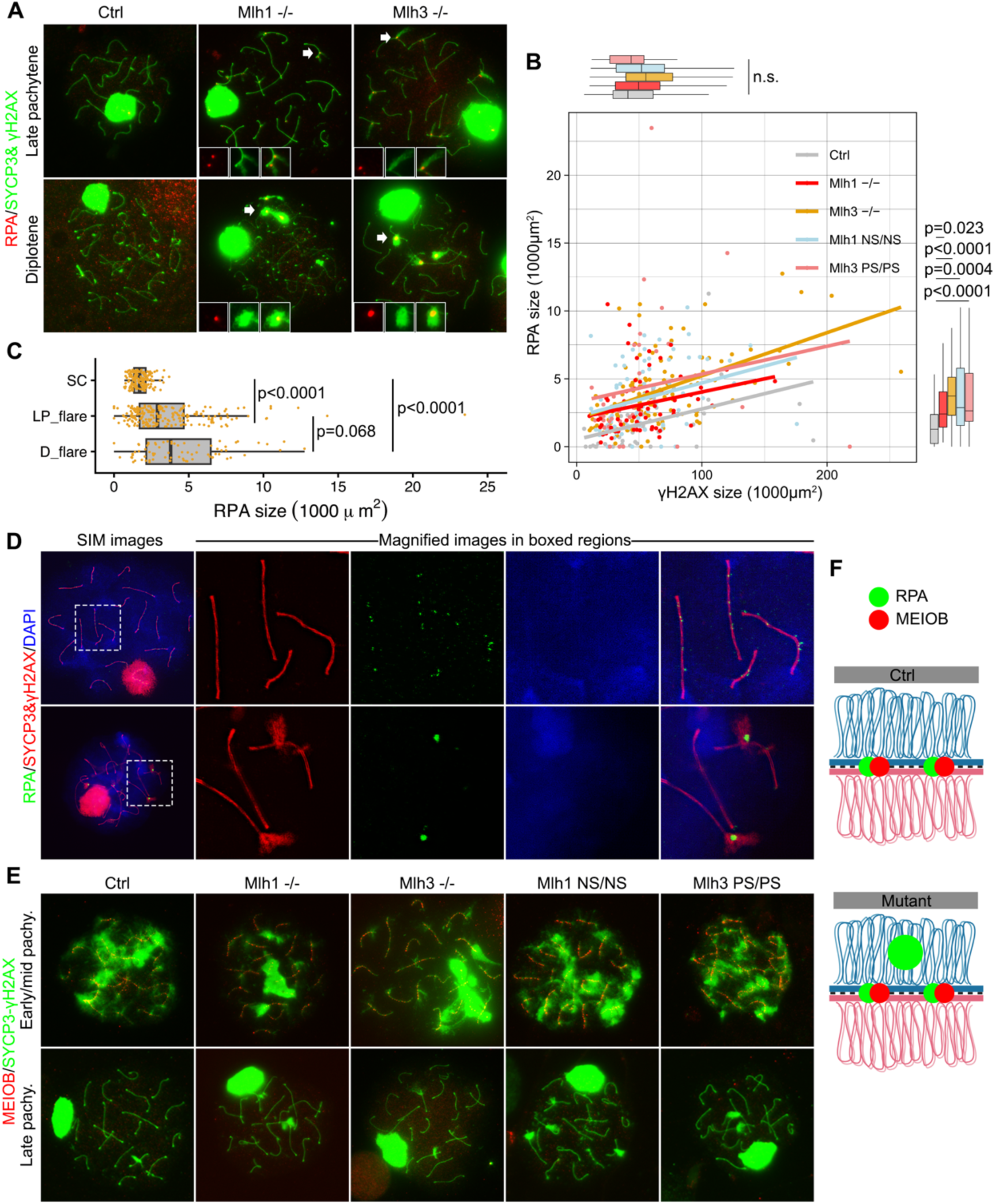
*Mlh1*/*3* deficiency induces unprogrammed DNA damage in spermatocytes. A) Representative images of RPA, γH2AX and SYCP3 staining of late pachytene and diplotene chromosome spreads from control (Ctrl) and mutant spermatocytes. Insets are magnifications of regions indicated by the white arrows, showing γH2AX flares. B) Comparison of RPA foci and γH2AX flare sizes. C) Distribution of RPA sizes on SC vs within flares. SC, synaptonemal complex; LP, late pachytene and D, diplotene. RPA sizes in LP_flare and D_flare were regrouped using data from mutant spermatocytes in panel B. D) SIM images of immunolabelled spreads of Ctrl (top row) and *Mlh1^-/-^* (bottom row) spermatocytes. The right-hand panels show a higher-magnification view of the dashed-box region. E) Immunostaining of SYCP3, γH2AX and MEIOB on Ctrl and mutant spermatocytes. MEIOB signal was absent in all 152 γH2AX flares analyzed from *Mlh1* and *Mlh3* mutant spermatocytes during late pachytene and diplotene stages. F) Schema illustrating the locations of RPA and MEIOB in Ctrl and *Mlh1/3* mutant spermatocytes. Data in B and C were analyzed using one-way ANOVA with Tukey’s *post hoc* test. For boxplots in B and C, black central line represents the median, and boxes and whiskers represent the 25th and 75th and 2.5th and 97.5th percentiles, respectively.

Super-resolution structured illumination microscopy (SIM) revealed that RPA on SC-associated foci and RPA within γH2AX flares differed markedly in both localization and size, supporting the presence of extensive ssDNA on chromosomal loops in *Mlh1/3* mutant spermatocytes (Fig. 4D). Interestingly, although the meiosis-specific ssDNA-binding protein MEIOB, which is involved in stabilization of recombination intermediates (42), was readily detected along the SC during early and mid-pachynema, it was absent from flares at later stages, including late pachynema (Fig. 4E). Together, these findings indicate that γH2AX flares in *Mlh1/3* mutant spermatocytes contain extensive stretches of ssDNA on chromosomal loops that arise through ongoing DNA processing and are distinct from canonical meiotic recombination intermediates (Fig. 4F).

### *Mlh1/3* deficient spermatocytes exhibit DNA hyper-resection and HR

The presence of enlarged RPA-positive regions within γH2AX flares, together with their progressive increase in size during meiotic progression (Fig. 4A–E), suggested that these sites may represent abnormal DNA end resection. During canonical DNA end resection, CtIP (CtBP-interacting protein) functions with the MRN complex (composed of MRE11, RAD50, and NBS1) to produce short stretches of ssDNA at DSB ends. Subsequently, downstream nucleases and helicases such as exonuclease 1 (EXO1), DNA2 (DNA replication helicase/nuclease 2) and BLM (Bloom syndrome protein), are recruited to elongate the 3′-ssDNA for HRR (43, 44). A recent study revealed that the MRN complex is involved in initiating and extending DNA end resection in mouse meiosis (41). In addition, previous studies in cancer cells showed that loss of *Mlh1* can lead to unrestrained DNA excision by EXO1 (45). We therefore examined whether DNA resection-associated proteins were present within γH2AX flares. Co-immunostaining revealed that approximately 50% of γH2AX flares in *Mlh1/3* mutant spermatocytes contained RAD50, CtIP, or BLM foci, whereas these proteins were rarely detected in control spermatocytes (Fig. 5A–F). RPA frequently colocalized with these resection factors within γH2AX flares (Fig. 5G), suggesting that active DNA end resection occurs at these sites. These findings support a model in which loss or impairment of *Mlh1/3* permits excessive DNA end processing in mutant spermatocytes, resulting in hyper-resection within loop regions that extend outward from the SC (Fig. 5H).

**Figure 5.**
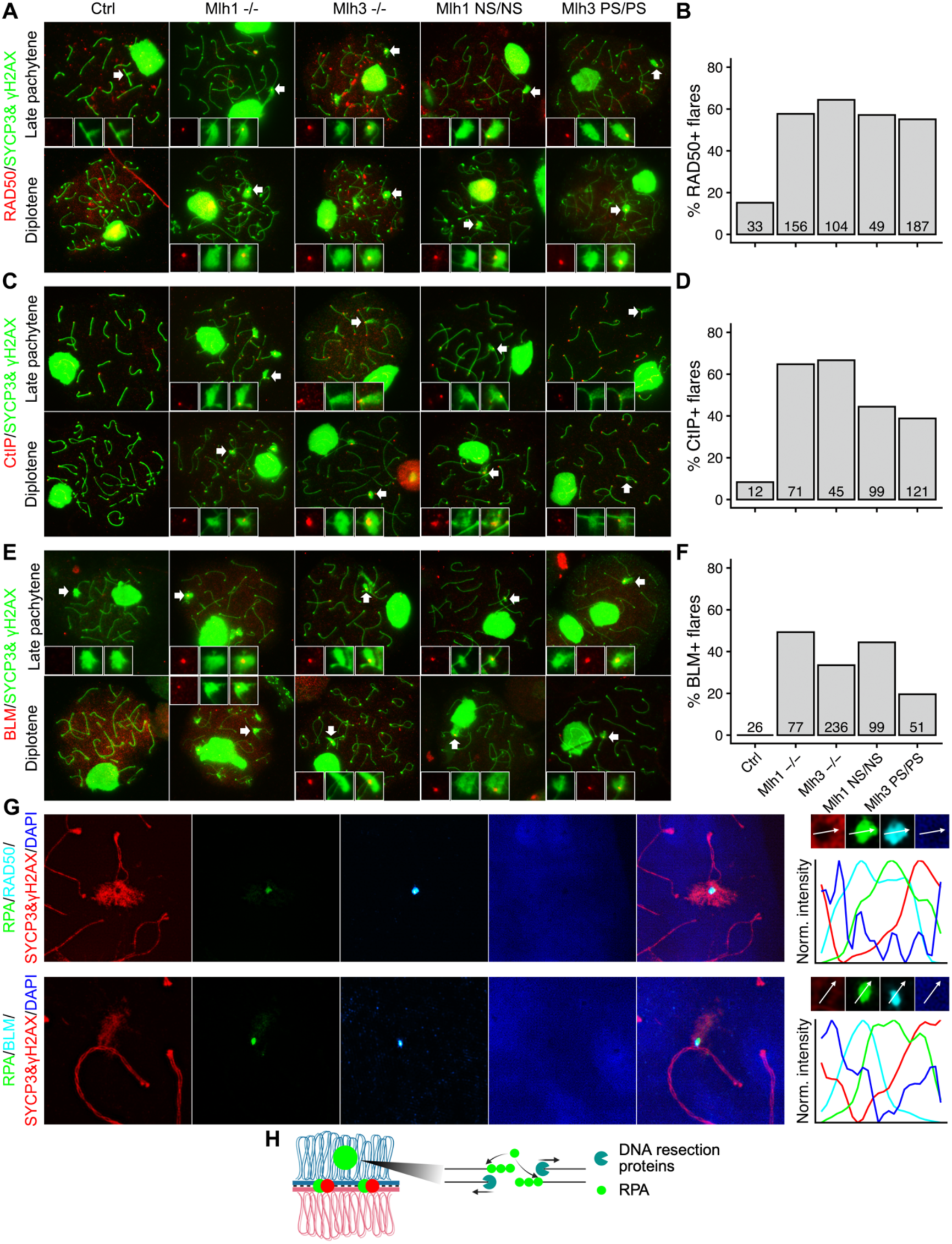
Extensive DNA end resection in late prophase I *Mlh1* and *Mlh3* deficient spermatocytes. A, C, E) Triple immunostaining of meiotic chromosome spreads for γH2AX, SYCP3 and either RAD50 (A), CtIP (C) or BLM (E). B, D, F) Fractions of γH2AX flares containing either RAD50 (B), CtIP (D), or BLM (F). G) SIM images of quadruple staining of RAD50 or BLM with RPA, γH2AX and SYCP3 in mutant spermatocytes. Right panel is the line-scan analysis of foci along the indicated line. Fluorescence intensities were independently normalized to a range of 0-1 for each channel. Line colors correspond to the respective fluorescence channel colors shown in the images. H) Proposed schema illustrating DNA resection in late stages of prophase I in *Mlh1/3* mutant spermatocytes. The magnified insets in A, C and E show the region of γH2AX flares, as indicated by white arrows. Number on each bar in B, D and F indicates the # of spermatocytes analyzed. Genotypes in A, C and E are at top of “A”.

To determine whether these resected regions initiate homologous recombination, we assessed RAD51/DMC1 localization within γH2AX flares (the anti-RAD51 Ab cross-reacts with both, but hereafter “RAD51” is used). In *Mlh3*^-/-^ spermatocytes, > 60% of flares contained RAD51, indicating active HR at these sites (Fig. 6A and B). By contrast, RAD51 was detected in only ∼20% of flares in the other mutants, a frequency comparable to controls (Fig. 6A and B). SIM imaging further confirmed colocalization of RAD51 and RPA within γH2AX flares (Fig. 6C). Together, these results indicate that γH2AX flares in *Mlh1/3* mutant spermatocytes are associated with extensive DNA end resection and, in some contexts, recruitment of RAD51, consistent with hyper-resection and activation of HR at these sites.

**Figure 6.**
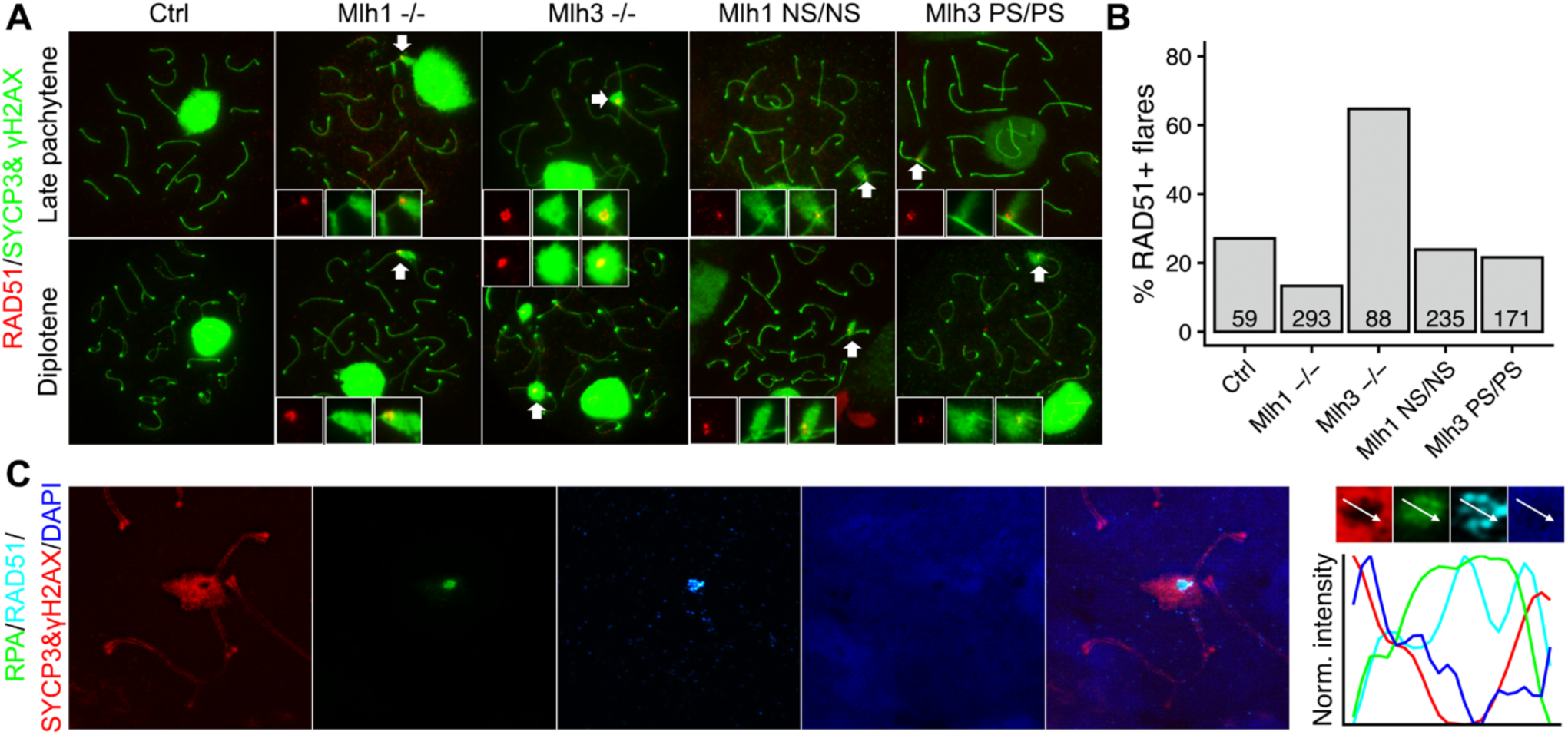
Evidence of homologous recombination repair in late prophase I *Mlh1* and *Mlh3* deficient spermatocytes. A) Representative images of spermatocyte chromosome spreads triply stained for RAD51, γH2AX and SYCP3. The insets are magnifications of γH2AX flares indicated by the white arrows. B) Quantification of γH2AX flares containing RAD51 foci. Numbers indicate the spermatocytes quantified. C) SIM images of quadruple staining for RAD51, RPA, γH2AX and SYCP3 in Mlh3^-/-^ spermatocytes. Right panel is the line-scan analysis of focus along the indicated line. Fluorescence intensities were independently normalized to a range of 0-1 for each channel. Line colors correspond to the respective fluorescence channel colors shown in the images.

**Figure 7.**
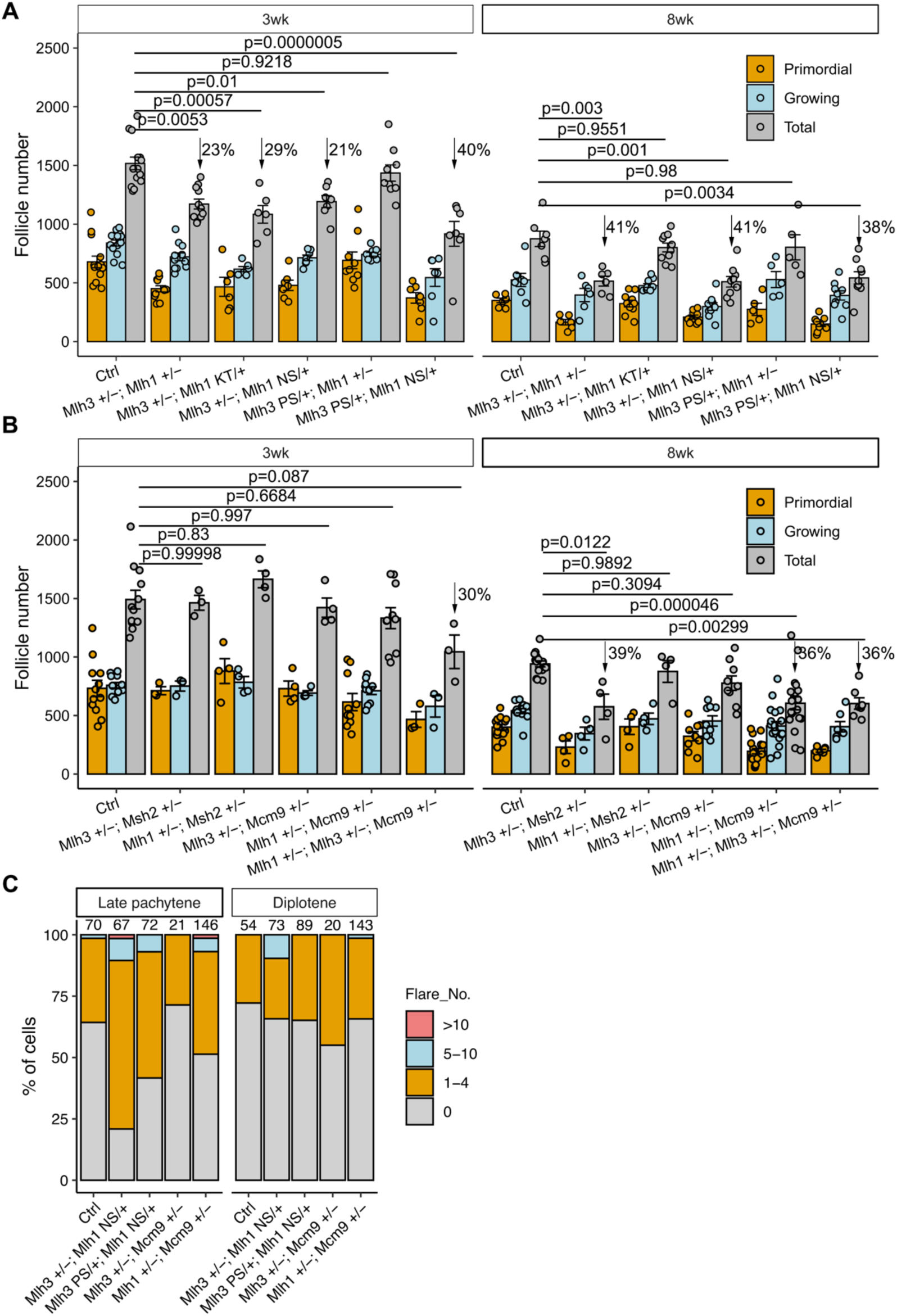
Oligogenic phenotypes of MMR mutants. A, B) Follicle number per ovary in 3wk- and 8wk-old mice having indicated genotypes. C) Quantification and distribution of gH2AX flares on autosomes in mutant spermatocytes. Number above each column indicates the spermatocytes quantified. Data in A and B were represented as the mean ± SEM and were analyzed using one-way ANOVA with Tukey’s *post hoc* test.

### Compound heterozygosity of *Mlh1* and *Mlh3* alleles decreases the ovarian reserve

A major challenge in human genetics is to identify the causative variant(s) responsible for an individual’s disease phenotype, especially in polygenic situations. Resequencing of patients may identify monoallelic variants in candidate genes, but causation is questionable if the allele is recessive (as with those studied here). However, such heterozygous alleles can contribute to a phenotype in the context of monoallelic disruptions in other genes in the same pathway, a phenomenon that has long been recognized and exploited in genetic studies of various developmental and biochemical processes. Genes underlying phenotypes with digenic or oligogenic inheritance are often enriched for shared biological functions, including common Gene Ontology annotations, subcellular localization patterns, protein–protein interactions, and co-expression relationships (46). Because *Mlh1* and *Mlh3* functionally and physically interact as components of the MutLγ complex, we hypothesized that combining monoallelic disruption of both genes might cause a phenotype that recapitulates homozygosity for either allele.

To test this, we generated several compound heterozygous combinations of knock-in and null alleles. We also include *Mlh1^K618T^*(KT) mice, which exhibit CO defects and POI (16). Male mice carrying various combinations of null and missense alleles, including *Mlh1*^+/-^ *Mlh3^+/-^* showed normal testicular architecture and sperm numbers comparable to WT and single heterozygote controls (Fig. S9). In contrast, females bearing most compound heterozygote combinations (*Mlh1^+/-^ Mlh3^+/-^*; *Mlh1^NS/+^ Mlh3^+/-^*; *Mlh1^KT/+^ Mlh3^+/-^* and *Mlh1^NS/+^ Mlh3^PS/+^)* had decreased follicle numbers (Fig. 8A). Interestingly, follicle loss in these compound heterozygotes was less severe than that observed in homozygous mutants, indicating that ovarian reserve maintenance is sensitive to the cumulative dosage of functional MutLγ. These findings support a dosage-dependent interaction between the two genes, in which partial impairment of both proteins is sufficient to compromise follicle maintenance, while complete loss of function results in a substantially more severe phenotype.

To determine whether follicle loss also occurs through interactions between *Mlh1/3* and other MMR genes, we generated compound heterozygotes involving *Mlh1*, *Mlh3*, *Mcm9*, and *Msh2*. Regarding *Mcm9*, aside from its well characterized role in HRR, there is a single report that it is involved in MMR based on its perturbation causing microsatellite instability, presence in a complex with other MMR proteins including MSH2 and MLH1, and dependent upon MSH2 for loading onto chromatin (47). *Mcm9* hypomorphic mice have reduced germ-cell numbers at birth due to impaired PGC proliferation, and *Mcm9*^−/−^ females are sterile, while males show markedly reduced sperm production and meiotic recombination defects (28, 48, 49). *Msh2*^−/−^ mice are fertile and have near normal sperm counts, though some seminiferous tubules lack germ cells (50). Spermatogenesis was unaffected in all compound heterozygote genotypes tested (*Mlh3^+/-^ Msh2^+/-^; Mlh1^+/-^ Msh2^+/-^; Mlh3^+/-^ Mcm9^+/-^* and *Mlh1^+/-^ Mcm9^+/-^*) (Fig. S9), and various compound mutants had little or no increase in γH2AX flares. Additionally, follicle numbers in 3-week-old females were no different than controls, indicating that there was no PGC loss. However, 8-week-old *Mlh1^+/-^ Mcm9^+/-^* females had fewer follicles vs controls (Fig.8B).

To explore whether more complex genotypes could underlie reproductive defects, we generated mice triply heterozygous for *Mlh1*, *Mlh3*, and *Mcm9* null alleles. Such males showed normal spermatogenesis, whereas *Mlh1*^+/−^ *Mlh3*^+/−^ *Mcm9*^−/−^ males exhibited smaller seminiferous tubules than those in *Mcm9*^−/−^ males (Fig. S9). In contrast to males, triply heterozygous females had fewer follicles than either *Mlh1*^+/−^ *Mlh3*^+/−^ or *Mlh1*^+/−^ *Mcm9*^+/−^ mice. These data provide an example of complex oligogenic effects on ovarian reserve maintenance.

## Discussion

Deciphering the consequences of genetic variants remains a major challenge in clinical genetics. Because pathogenicity algorithms can have substantial false positive rates, we combined them with structure-based analyses to prioritize variants for the labor-intensive process of functional evaluation in mice (51)(52). This approach was validated in that 4 of the 5 models we generated exhibited reproductive defects. Overall, our studies show that partial loss-of-function *Mlh1/3* variants, as well as compound heterozygosity for null alleles, can compromise ovarian reserve maintenance without causing overt infertility. Additionally, the infertility of *Mlh1^G98S^* founder mice is a potential example of a dominant-negative allele that can provide precedence for clinical genetics. Structural analyses suggest that p.G98S does not abolish ATP/Mg²⁺ binding but instead perturbs the MLH1 ATPase domain in a manner likely to impair ATP-dependent conformational switching. Because MLH1 functions in a heteromeric complex, a variant protein that retains ligand binding and complex assembly but which is defective in ATPase function could potentially produce a dominant-negative effect, as in the case of an ATPase defective MutL in *E. coli* (59).

Studies on POI patients and mouse models have highlighted that defects in meiotic DSB formation and repair, chromosomal synapsis, and recombination, can compromise fertility and the ovarian reserve (3, 53). Although MutLγ is essential for ∼90-95% of all COs in mice, our findings reveal that ovarian reserve depletion can occur independently of detectable CO defects. *Mlh3^PS/PS^* and *Mlh1^NS/NS^* females exhibited follicle loss despite normal CO levels, whereas *Mlh1^KA/KA^* mice displayed both follicle depletion and reduced CO formation. Thus, the canonical CO-promoting role of MutLγ can be separated genetically from an additional function in oocyte survival.

An indication of this function was that all three mutants exhibited persistent autosomal γH2AX flares in late meiotic prophase. These flares were associated with unusually large RPA-positive regions and the DNA end-processing factors CtIP, RAD50, and BLM, suggesting persistent or excessive DNA resection, a process that is normally carried out by the MRN complex in processing SPO11-induced DSBs (41). The progressive enlargement of RPA-positive regions from late pachynema to diplonema further suggests that resection continues beyond the stage during which repair of programmed DSBs is normally completed, although it is unclear if these regions are related to sites of meiotic DSBs (see below). A similar phenomenon has been described in cancer cells, where MLH1 deficiency permits excessive EXO1-mediated DNA excision, leading to increased ssDNA formation, secondary DNA damage intermediates, and subsequent activation of the cGAS-STING pathway (45). Additionally, MLH1 was found to restrain DNA2 nuclease activity at stressed replication forks in BRCA2-deficient cells, enabling their survival (54). Although the nuclease(s) responsible for the meiotic molecular phenotype remain unknown, these findings raise the possibility that MutLγ normally limits resection or promotes termination of repair reactions during late prophase I.

In considering the nature of the repair response at these lesions, it appears to be distinct from a late-prophase response to exogenous DNA damage. Although NHEJ can be reactivated in late spermatocytes after irradiation (55), autosomal γH2AX flares in *Mlh1/3* mutants lacked the NHEJ marker 53BP1 and instead retained RPA and other resection factors, consistent with continued HR-type processing. The increased frequency of RAD51-positive flares in *Mlh3*-null spermatocytes may similarly reflect prolonged attempts to repair extensively resected DNA, consistent with delayed RAD51-focus resolution reported in *Mlh3*-deficient somatic cells (56). Because synapsis appears normal and programmed meiotic DSB repair is largely completed on schedule, we propose that these late DNA-processing defects activate ATM/ATR-dependent signaling and the CHK2-mediated oocyte quality-control pathway (20), leading to elimination of a subset of oocytes and depletion of the ovarian reserve.

The origin of the DNA lesions that undergo hyper-resection is unclear. Several lines of evidence argue against these structures being unrepaired programmed DSBs: 1) synapsis and CO levels were normal in *Mlh1^NS/NS^* and *Mlh3^PS/PS^*spermatocytes; 2) RAD51 foci along chromosome axes were not increased; 3) γH2AX flares extended from the synaptonemal complex into chromatin loop domains; and 4) the meiosis-specific ssDNA-binding protein MEIOB was absent from these structures. These findings indicate that the flares are distinct from canonical meiotic recombination intermediates. One possibility is that they originate from spontaneous DNA breaks within chromatin loops that are normally repaired or protected from excessive resection. Alternatively, in the absence of normal MutLγ activity, a subset of recombination intermediates may fail to be properly resolved and subsequently undergo aberrant nucleolytic processing during late prophase I, ultimately generating the extensive ssDNA tracts. Providing indirect support for this idea is that MutLγ was found to interact genetically and physically with the RecA-related Dmc1 recombinase in yeast to promote crossover formation (57).

Whole exome sequencing has facilitated the discovery of genetic mutants underlying monogenic disorders. Incomplete penetrance and variable expressivity suggest that additional genetic lesions shape clinical outcomes. Clinical studies have documented digenic inheritance in POI (58). Here, we uncovered evidence that *MLH1* and *MLH3* variants constitute such a digenic underpinning. Germline mutations in MMR-related genes often result in MSI and Lynch syndrome (LS, also known as hereditary nonpolyposis colorectal cancer). In one pedigree study, co-segregation of a missense variant in *MLH3* and a frameshift mutation in *MLH1* were identified in a LS patient (59). Thus, heterogeneous digenic pathogenic variants represent potentially important contributors to diseases such as POI and should be considered in genetic analyses. The MLH1-MCM9 interaction is also informative. Although MCM9 is known for its role in HR and germ cell maintenance (28, 49, 60), it has also been implicated in MMR through interactions with canonical MMR proteins, including MLH1 (47). Supporting this link, HORMAD1 interacts with the MCM8-MCM9 complex and impairs MLH1 chromatin loading, functionally linking MCM9-associated DNA repair pathways with MMR components (61). These overlapping surveillance mechanisms explain why combined heterozygosity can disrupt the ovarian reserve. Collectively, our findings reinforce the hypothesis that subtle defects across interacting repair pathways cumulatively compromise meiotic genome integrity and drive POI.

Collectively, our findings support a model in which MutLγ serves dual functions during meiosis. In addition to its established role in CO formation, MutLγ protects meiotic chromosomes from excessive DNA end processing during late prophase I. Disruption of this genome-protective activity results in progressive hyper-resection, activation of CHK2-dependent quality control pathways, and depletion of the ovarian reserve. This previously unrecognized function of MLH1 and MLH3 provides a mechanistic explanation for how hypomorphic variants can contribute to reproductive aging, subfertility, and POI even in the absence of overt CO defects.

## Acknowledgements

This work was supported by grants P50HD096723, R01HD082568 and P50HD104454 from the National Institute of Child Health and Human Development. We thank R. Munroe and C. Abratte of the Cornell Transgenic Core Facility for generating knock-in mice. Additionally, we thank Mary Ann Handel, Paula Cohen and Eric Alani for feedback on the manuscript, P. Cohen for Stra8-Cre mice (who in turn received them from R. Weiss at Cornell), and members of the Cohen and Schimenti labs for continued input on phenotype analysis, technical advice and reagents. We also would like to thank R. Weiss, R. Freire, J. Wang, and M.A. Handel for providing antibodies.

## MATERIALS and METHODS

### Prioritization of missense variants

Missense variants with a SNP rsID in human MLH1 and MLH3 were retrieved from gnomAD 2.1.1 database (gnomeAD.broadinstitute.org). Variant pathogenicity was assessed using dbNSFP v4.1a (62), which is integrated into the Ensembl Variant Effect Predictor (VEP) (63). Prediction outcomes from SIFT (64), PolyPhen-2 (65), REVEL (66), PROVEAN (67) and CADD (68) were integrated to prioritize potentially deleterious variants. The allele frequencies were obtained from gnomAD. To further evaluate the functional consequences of the prioritized variants, AlphaFold-predicted 3D structures of human MLH1 and MLH3 were used to assess the effects of missense substitutions on protein stability with the mCSM (21) algorithm.

### Structural modeling and ATP pocket analysis

AlphaFold 3 (24) was used to predict the structure of the human MLH1 ATPase domain (residues 1-347) in an ATP- and Mg²⁺-bound state. Models were generated for WT MLH1 and the p.N64S, p.G67R, and p.G98S variants using identical input settings with a random seed of 10. The top-ranked model for each construct was selected for downstream structural analyses. Binding-pocket prediction was performed using P2Rank (v2.5.1) (69). The top-ranked pockets identified in each WT and mutant structure were compared to assess alterations in ATPase-pocket organization. ATPase-pocket geometry was quantified by calculating the Euclidean distance between the Cα atom of each ATPase residue and the center of the top-ranked P2Rank-predicted pocket. To quantify pocket conservation relative to WT, Jaccard similarity coefficient was calculated by:

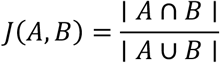

where *A* and *B*represent the pocket-residue sets of the WT and variant models, respectively.

Structural differences between WT and variant ATP-Mg-bound models were evaluated using ProDy (v2.4.1) (70). Cα atoms from each variant were aligned to the WT structure using the ProDy implementation of least-squares structural superposition. Per-residue displacement was calculated as the Euclidean distance between aligned WT and variant Cα coordinates. pLDDT scores were extracted from the B-factor field of AlphaFold 3-generated PDB files and used as a residue-level confidence metric. Residue-wise displacement profiles were generated to identify localized regions of structural remodeling. Missense3D (v2) was used to predict structural changes caused by missense variants (71).

### Mice

All animal usage was approved by Cornell University’s Institutional Animal Care and Use Committee, under protocol 2004-0038 to J.C.S.

The CRISPR/Cas9 genome editing technology was applied to generate humanized/ knock-in and *Mlh1/3* knockout mice, as previously described (22, 51, 72). sgRNAs and ssODNs are listed in Table S1. B6.Cg-*Msh2*^tm2.1Rak^/J was purchased from The Jackson Laboratory (JAX:016231), and the Stra8-Cre driver line (73) was obtained from Paula Cohen. The *Chk2* knockout strain used was *Chk2^tm1Mak^*as previously described (74). *Mcm9* gene trap mice (AWO655 clone) were described previously (28). *Mlh1^K618T^* and *Mlh1^K618A^* mice were constructed previously in our lab (16). The primers used for genotyping are listed in Table S1. To distinguish WT and mutant alleles, PCR amplicons were analyzed via Sanger sequencing or restriction enzyme digestion. For identification of *Mlh3^P1262S^* and *Mlh3* null mutants, amplicons were digested with restriction enzyme *Bsl*I (NEB, R0555S) at 37°C for 1 hour and products were analyzed on high percentage agarose gels. Digestion of WT products yield 135bp and 153bp bands, whereas the *Mlh3^P1262S^* and *Mlh3* null amplicons remained undigested, yielding single bands of 288bp and 283bp, respectively.

### Fertility test, sperm count, histology analysis and follicle quantification

Fertility of mice was assessed by housing mutant mice with opposite-sex WT animals and recording litter sizes over time. Sperm counts, histological analyses and ovarian follicle quantification were performed as previously described (72). For follicle quantification, every fifth section was examined for the presence of the following classes of oocytes/follicles: primordial, primary, secondary, preantral, and antral. For quantitative analyses, they were grouped into two classes: primordial and growing (primary through antral).

### Meiotic spread preparation and immunostaining

Meiotic prophase I surface spreads were prepared as described previously (75). Briefly, the testes were removed and decapsulated in hypotonic sucrose extraction buffer and left on ice for 1h. Tubules were chopped on glass depression slides in a bubble of 0.03% sucrose and added to slides coated in 1% paraformaldehyde (PFA). The slides were slow dried and subjected to immunofluorescence analyzes.

Prophase I stages were determined based on SYCP3 staining patterns (4). For the staining, the slides were incubated for 1h at RT with sterile filtered antibody dilution buffer (ADB) containing 10 ml normal goat serum, 3 g bovine serum albumin, 50 μL Triton X-100 and 990 mL PBS, and subsequently incubated overnight with ADB diluted primary antibodies list in Table S2 in a humidified chamber at 4 °C. After washing in ADB, the slides were incubated at 37 °C for 30 min with a 1:1000 dilution of secondary antibody goat anti-rabbit IgG 594 (Invitrogen, A11012) and goat anti-mouse IgG 488 (Invitrogen, A11001), then incubated at 37 °C for 5 min with 500 ng/mL of DAPI and mounted as described above.

TUNEL staining was performed using the DeadEnd™ Fluorometric TUNEL System (Promega, G3250) following manufacturer’s instructions. TUNEL and PH3 double immunostaining was performed as previously described (16).

### Diakinesis/ Metaphase chromosome preparations

This procedure was performed as described (16). Briefly, seminiferous tubules were treated by hypotonic solution (1% sodium citrate) for 30 min and minced carefully for 15 min. The cell suspension was centrifuged, and the supernatant was removed. The cells were then fixed three times in methanol: glacial acetic acid (3:1) and dropped onto slides. After drying, the chromosomes were stained with Giemsa solution prepared from Giemsa powder (Sigma, G4507).

### PGCLC differentiation

To construct missense alleles in ESCs, pSpCas9(BB)-2A-PuroR (PX459) V2.0 vector (Addgene, #62988) harboring gRNA was co-transfected along with the corresponding ssODN. After puromycin selection for 48 hours, the single colony was expanded and genotyped. The sequences of all the gRNAs, ssODN and primers were the same as those used to generate the corresponding mouse models and are provided in Table S1. Cell lines with frameshift mutations in *Mlh1* or *Mlh3* were also expanded to serve as knockout allele for comparison. PGCLC differentiation and analysis were performed as described previously (31).

### Image process and data statistics

Images were captured by an Olympus BX51 microscope and Olympus cellSens standard software or Zeiss Axiophot Z1 microscope attached to a cooled charge-coupled device (CCD) Black and White Camera (Zeiss McM) and pseudo-colored using ZEN 2 software (Carl Zeiss AG, Oberkochen. Germany). Higher resolution images were acquired using a Zeiss ELYRA 3D-Structured Illumination Super Resolution Microscopy (3D-SIM) with ZEN Black software (Carl Zeiss AG, Oberkochen. Germany). Cropping, color, and contrast adjustments were made with Adobe Photoshop CC 2025, using identical background adjustments for all images.

Fluorescence intensity was measured by ImageJ. All data were expressed as mean ± SEM. Statistical tests were carried out using a paired or unpaired student’s *t-test* or one-way analysis of variance (ANOVA) followed by the Tukey’s *post hoc* test with R. Graph generation was performed using R. The schematic diagrams were created using BioRender. Protein structures were visualized using PyMOL (v.3.1.8, Schrödinger, LLC).

## Competing interests

The authors declare no competing or financial interests.

## Notes

### Competing Interest Statement

The authors have declared no competing interest.

