## Supplemental Figures 1-9 for "Modeling of human *MLH1* and *MLH3* variants in mice reveals novel functions of MutLγ in meiotic DNA repair and the ovarian reserve"

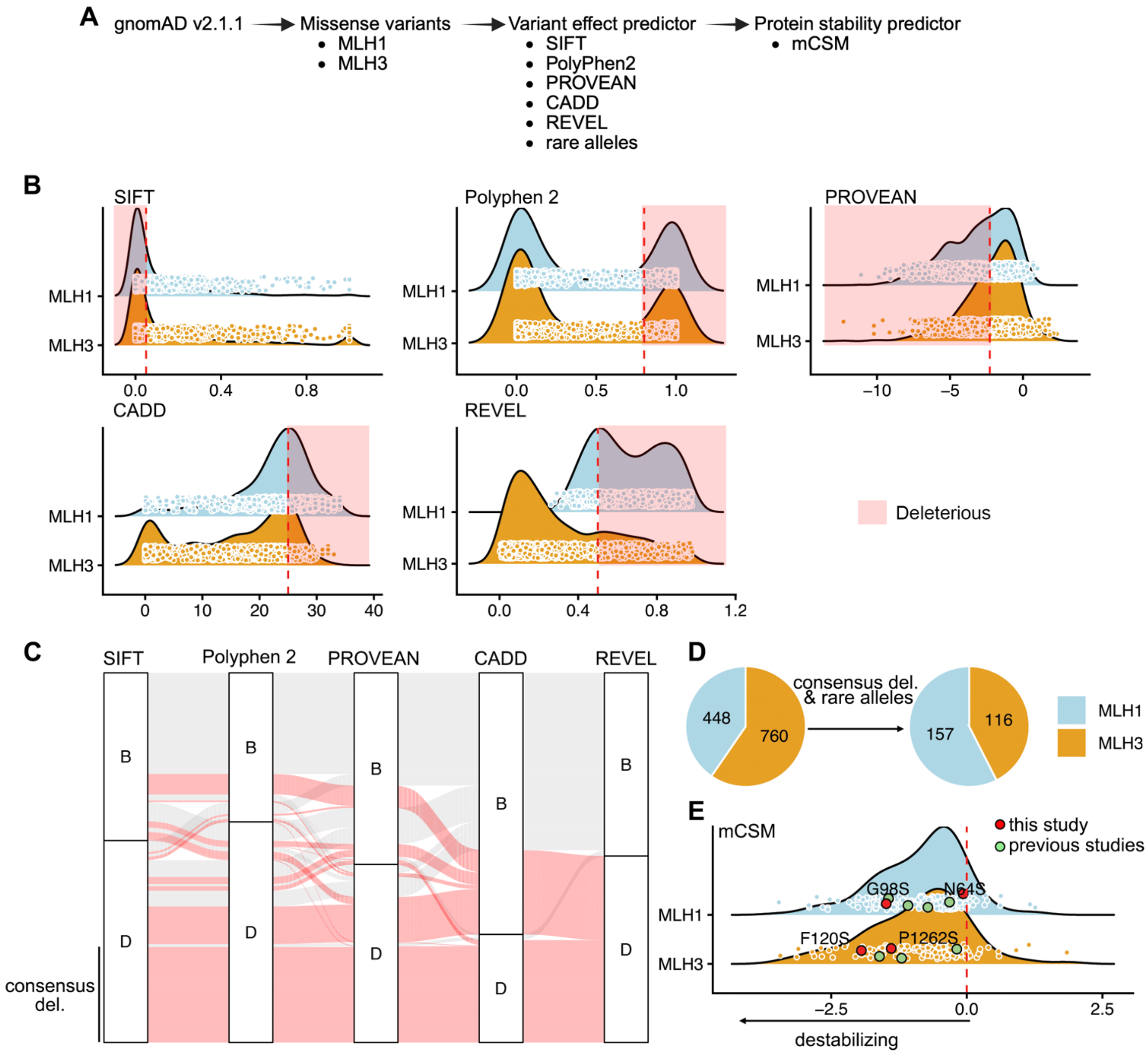


**Figure S1. Prioritization and functional prediction of rare deleterious missense variants in *MLH1* and *MLH3*.** A) Pipeline of variant prioritization. B) Distribution of prediction scores by 5 different *in silico* algorithms. Red dashed lines indicate the deleteriousness thresholds for each predictor, and the pink rectangle highlight the score ranges classified as deleterious. C) Inconsistent prediction outcomes across algorithms and consensus identification of deleterious variants were prioritized for further study. D) Distribution of rare missense variants predicted to be deleterious in MLH1 and MLH3. E) Predicted effects of deleterious missense variants on protein stability, assessed using the mCSM algorithm (n = 157 for MLH1 and n = 116 for MLH3). Positive ΔΔG values indicate increased protein stability, whereas negative ΔΔG values indicate destabilization. Missense variants selected for mouse model generation are highlighted with larger symbols. Variants previously investigated (Refs 16 and 22) and are indicated by green dots, whereas variants generated in the present study are indicated by red dots. Previously modeled alleles (green dots) are MLH3 p.R93G, MLH3 p.R1230H, MLH3 p.A1394T, MLH1 p.K618E, MLH1 p.V384D, MLH1 p.K618T, and MLH1 p.E268G.

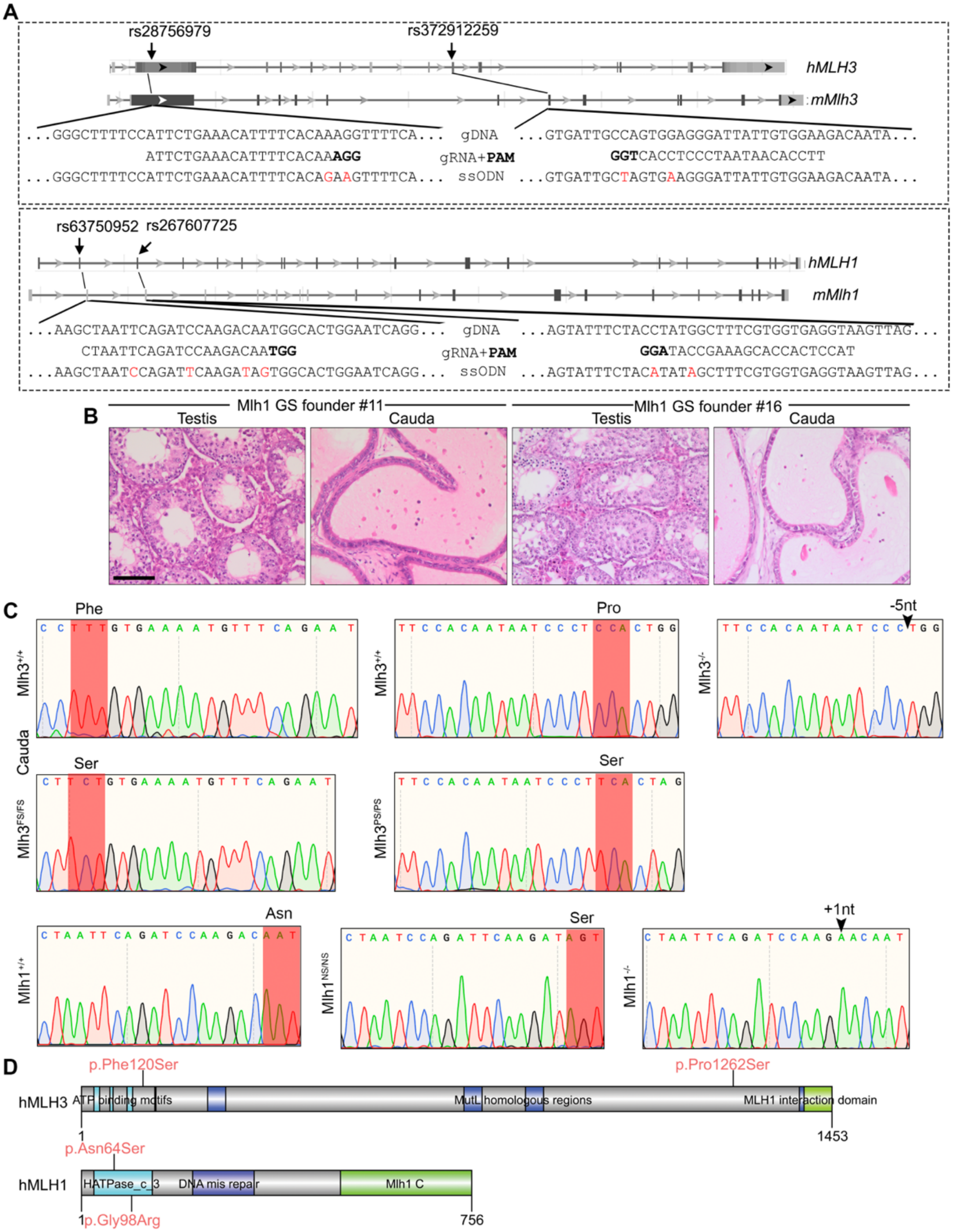


**Figure S2. Modeling *MLH1/3* missense mutations in mice.** A) Diagram of CRISPR-Cas9 genome editing strategy to introduce the p. Phe120Ser and p. Pro1262Ser amino acid substitutions in mouse *Mlh3*, and the p. Asn64Ser and p. Gly98Ser amino acid substitutions in mouse *Mlh1*. The human *MLH3* p. Phe120Ser and p. Pro1262Ser variants encoded by SNP rs28756979 and rs372912259 are in exon 2 and 8, respectively. The human *MLH1* p. Asn64Ser and p. Gly98Ser variants encoded by SNP rs63750952 and rs267607725 are in exons 2 and 3, respectively. The bold fonts indicate the PAM and the red fonts represent the introduced substitutions of nucleotide. B) Representative histological images of testis and cauda epididymides from *Mlh1* GS founder mice. The Mlh1 p.G98S allele in these founder mice was identified using TA cloning. Scale bar = 100 μm. C) Representative Sanger sequencing chromatograms from indicated mutant mice. The relevant codon bases were labeled above and were highlighted by red box. Black arrowhead above the chromatogram indicates the deletion or insertion. D) Schematic of domain structure of the human MLH3 and MLH1 proteins and the variant location.

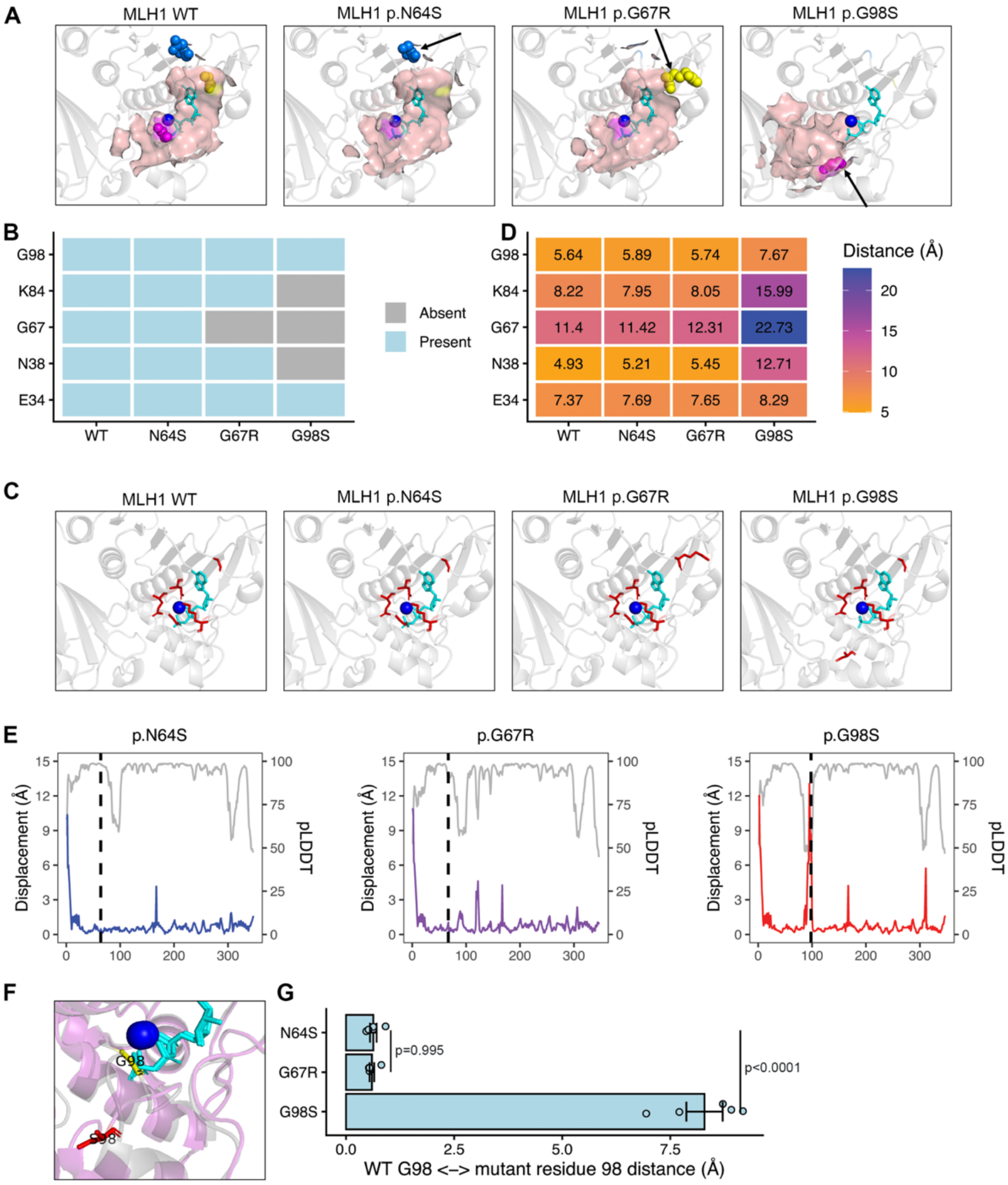


**Figure S3. Structural effects of MLH1 ATPase-domain variants.** A) Predicted ATP-$Mg^{2+}$ binding pockets identified by P2Rank in AlphaFold3 models of indicated MLH1 variants. Protein backbones are shown as a gray cartoon, and the predicted pocket surfaces are shown in pink. ATP is depicted as cyan sticks and $Mg^{2+}$ as a blue sphere. Variant residues are highlighted as spheres and indicated by arrows. B) Presence or absence of key ATPase-pocket residues within the top-ranked P2Rank pocket predicted for WT and mutant MLH1 proteins. C) Localization of key ATPase-pocket residues in the MLH1 ATPase domain. Protein backbones are shown as gray cartoons, ATPase-pocket residues are shown as red sticks, ATP is displayed as cyan sticks, and $Mg^{2+}$ is shown as a blue sphere. D) Distances between key ATPase-pocket residues and the center of the predicted pocket in WT and mutant MLH1 models. Distances were calculated using residue Cα coordinates and the pocket-center coordinates. E) Residue-wise structural displacement between WT and mutant MLH1 models following structural superposition. Colored traces indicate Cα displacement relative to WT (left y-axis), whereas gray traces indicate AlphaFold3 confidence scores (pLDDT; right y-axis). Vertical dashed lines indicate mutation sites. F) Structural superposition of WT model (grey) and the merged G98S mutant (purple). Residues G98 (yellow) and S98 (red) are highlighted as sticks. ATP is depicted as cyan sticks and $Mg^{2+}$ as a blue sphere. G) Distance between residue 98 in the top-ranked WT model and the corresponding residue in structurally aligned mutant models. Each dot represents an independent AlphaFold3 prediction. Data are represented as the mean± SEM and were analyzed using one-way ANOVA with Tukey's *post hoc* test.

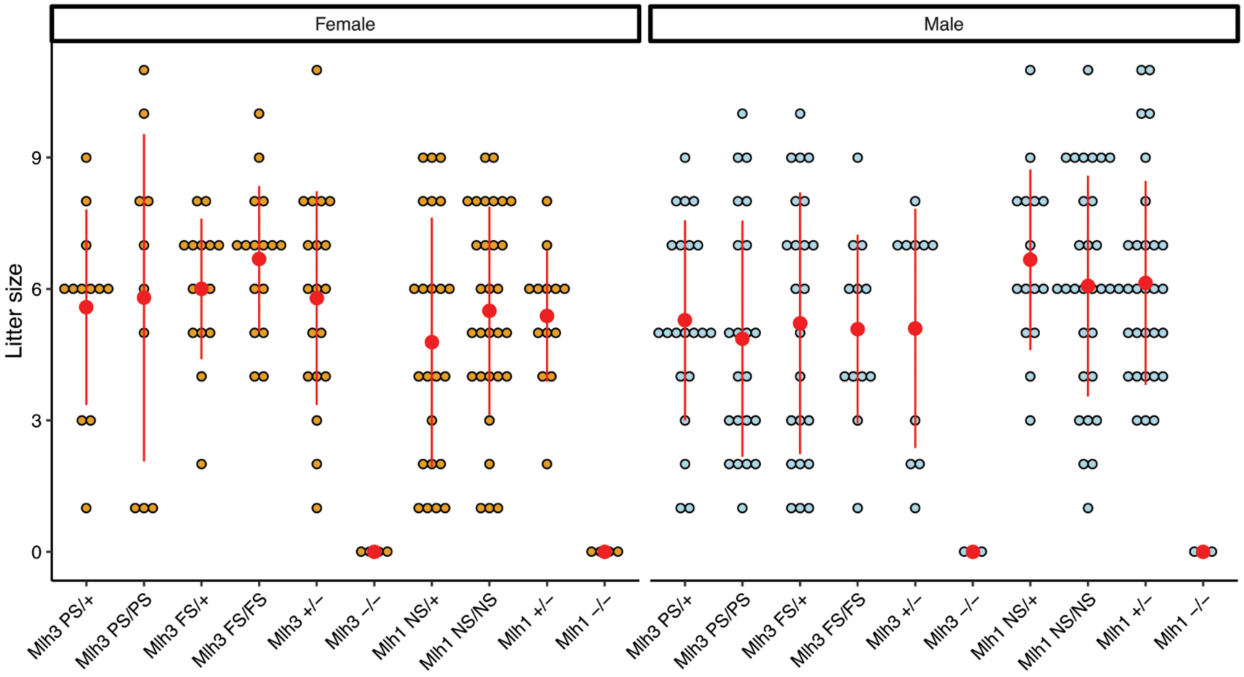


**Figure S4. Litter sizes from matings of mutant mice to opposite sex WT partners.**

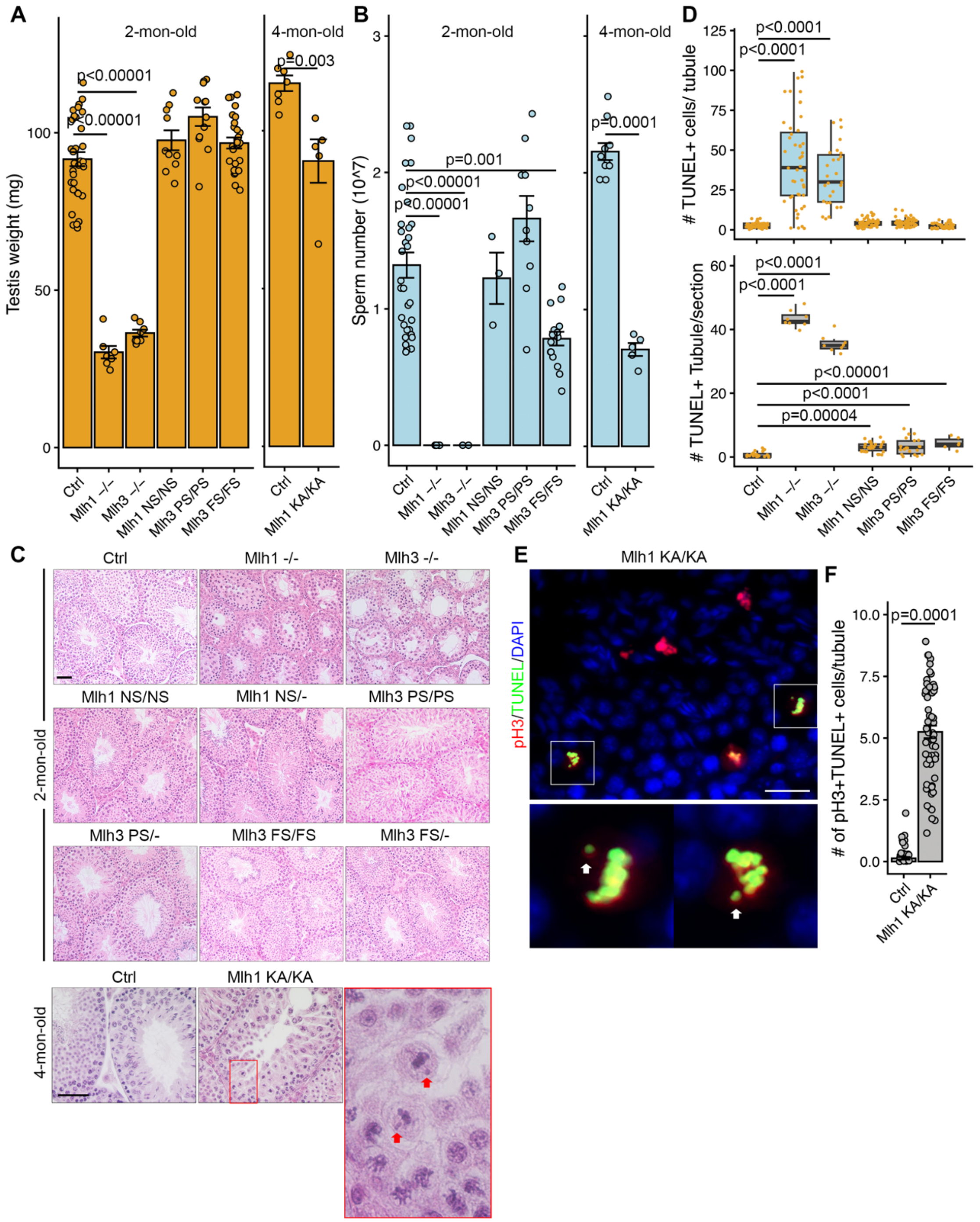


**Figure S5. Phenotyping of *Mlh1* and *Mlh3* mutant male mice.** A) Testis weights. B) Sperm counts. C) Representative histological images of testis from control (Ctrl) and mutant mice. Scale bars = 50 µm. D) Quantification of TUNEL$^+$ cells in each tubule. E) Number of tubules containing more than 4 TUNEL$^+$ cells in each section. This number was chosen as control tubule section had >4. F) Dual staining of pH3 and TUNEL on 4-month-old KA/KA testis. Scale bars = 50 µm. G) Number of pH3 and TUNEL double positive cells in each tubule. The arrows in higher-magnification view of the box region in C and E indicate the lagging chromosomes. Data in A, B and F are represented as the mean± SEM. Data in A, B and D were analyzed using one-way ANOVA with Tukey's *post hoc* test. Data in F were analyzed using a two-tailed unpaired *t* test. In D, black central line represents the median, and boxes and whiskers represent the 25th and 75th and 2.5th and 97.5th percentiles, respectively.

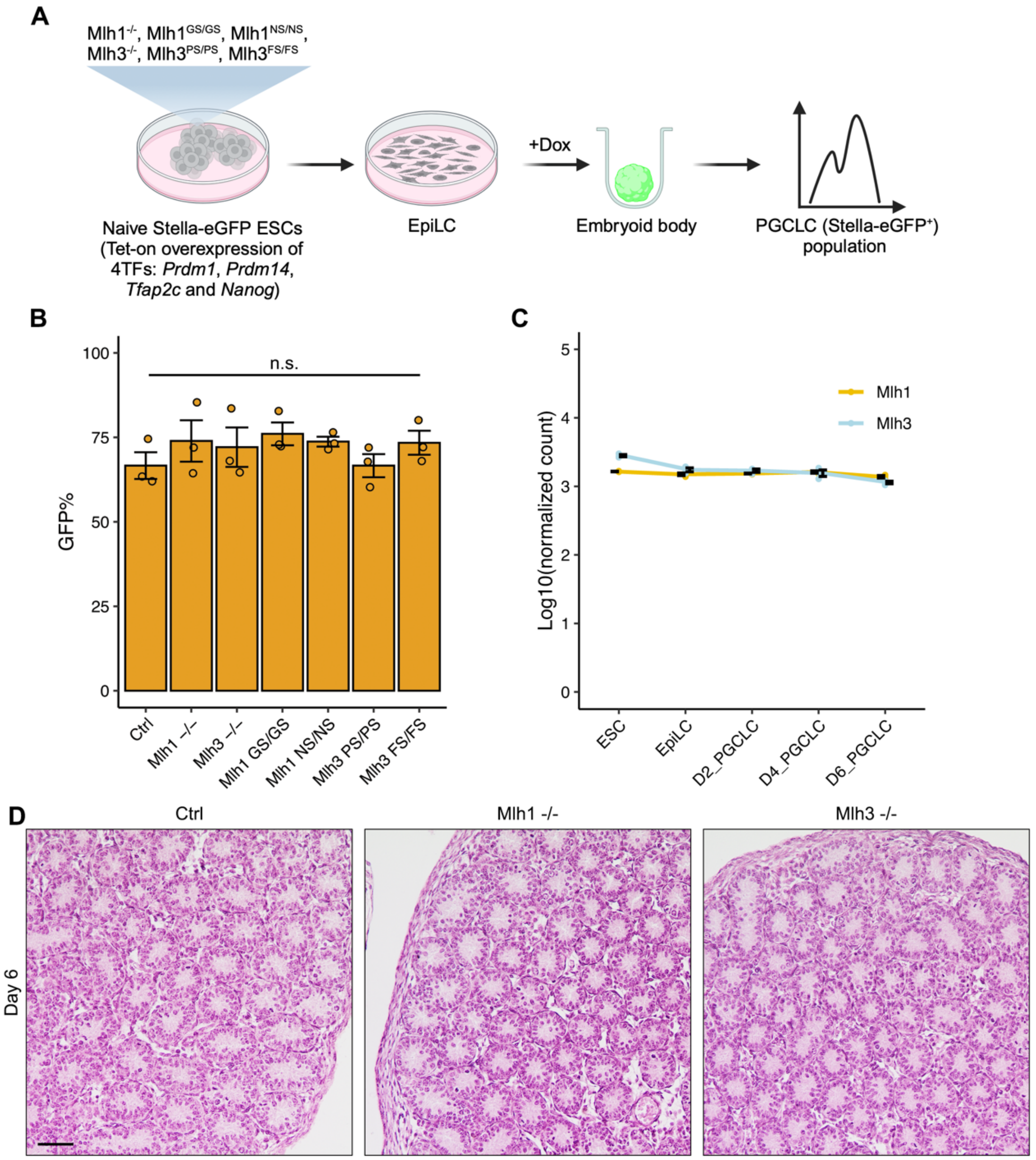


**Figure S6. *Mlh1* and *Mlh3* are dispensable for PGCLC specification *in vitro* and spermatogonia development *in vivo*.** A) Scheme of *in vitro* embryonic stem cell differentiation. *Stella-eGFP* was used to indicate the PGCLC population in embryoid bodies. B) Quantification of GFP cells (PGCLCs) at day 6 differentiated from *Stella-eGFP* ESCs with point mutants (*Mlh1*$^{GS/GS}$, *Mlh1*$^{NS/NS}$, *Mlh3*$^{PS/PS}$, and *Mlh3*$^{FS/FS}$) or frameshifts (*Mlh1*$^{-/-}$ and *Mlh3*$^{-/-}$). The unedited parental cells serve as control (Ctrl). *Mlh1*$^{-/-}$ and *Mlh3*$^{-/-}$ ESCs harbor frameshift mutations distinct from those in the mouse models. C) Gene expression levels of *Mlh1* and *Mlh3* in ESC, EpiLC and PGCLC at days 2/4/6 (D2/4/6). D) Representative histological images of testis from 6-day-old Ctrl and mutant mice. Scale bar = 50 µm. Data in B and C are represented as the mean± SEM. Data in B were analyzed using one-way ANOVA with Tukey's *post hoc* test.

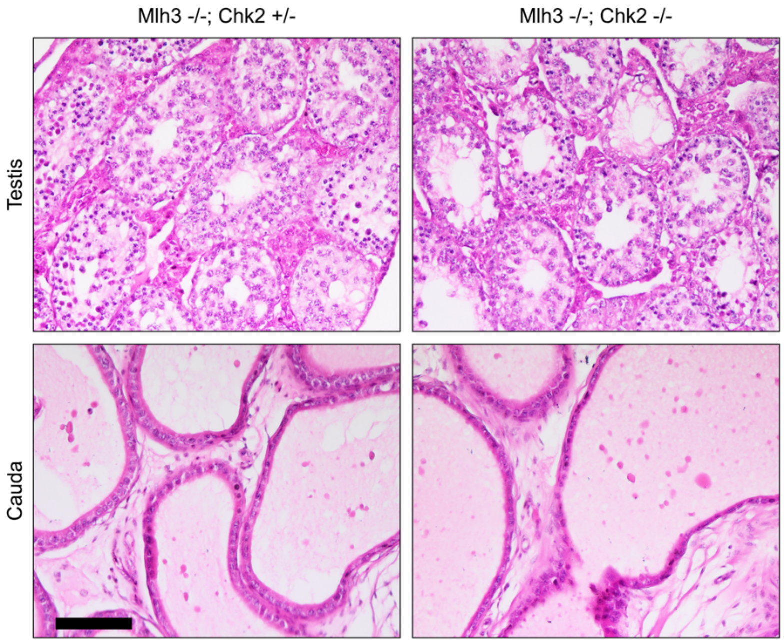


**Figure S7. Representative histological images of testis and cauda epididymides from 4-month-old mutant mice.** Scale bar = 100 μm.

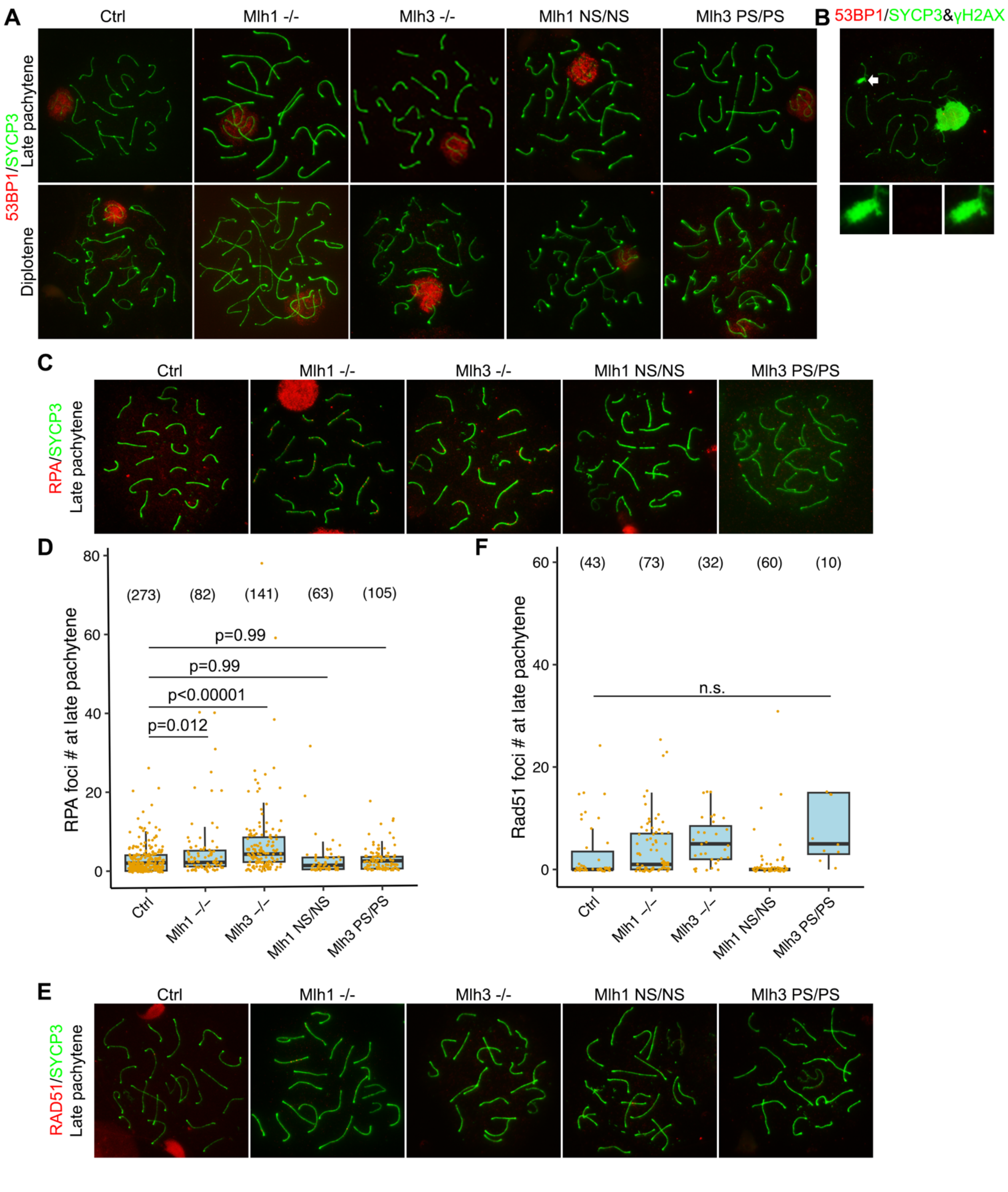


**Figure S8. Characterization of 53BP1, RPA and RAD51 in spermatocytes.** Representative images of mutant spermatocyte chromosome spreads immunolabelled for A) 53BP1; B) 53BP1 and γH2AX, with insets showing magnifications of γH2AX flares indicated by white arrows; and C) RPA, quantified in (D). E) Immunolabeling of RAD51 in late pachynema and quantified in (F). n.s., no significant difference. Data in D and F were analyzed using one-way ANOVA with Tukey's *post hoc* test. Numbers in brackets in D and F indicate the number of spermatocytes quantified.

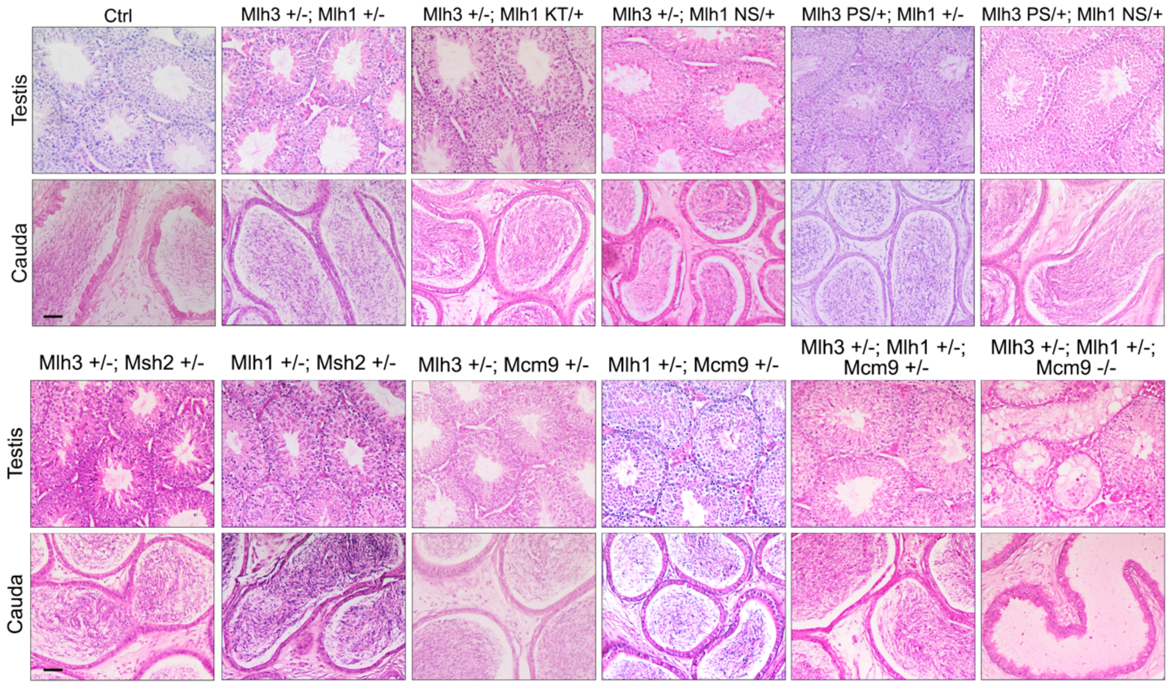


**Figure S9. Representative histological images of testis and cauda epididymides from 8wk-old mice.** Scale bar = 50 µm.
