## Supplementary text for "Modeling of human *MLH1* and *MLH3* variants in mice reveals novel functions of MutLγ in meiotic DNA repair and the ovarian reserve"

### *In silico* structural analysis of the MLH1 p.G98S ATPase-domain variant.

To investigate why an *Mlh1 p.G98S* knock-in mouse line could not be established, we performed comparative structural modeling of the MLH1 ATPase domain. Gly98 lies within the conserved GFRGE(A/G)L motif and packs near Gly101 adjacent to the ADP pyrophosphate moiety (1), placing it in a region likely to influence nucleotide-dependent conformational changes. Using AlphaFold 3, we modeled ATP/$Mg^{2+}$-bound human MLH1 structures for WT, p.G98S, p.N64S, and p.G67R; the latter has previously been shown to impair mismatch repair and meiotic crossover formation (2). P2Rank (3) pocket analysis identified G67 and G98, but not N64, as residues contributing to the predicted ATP-binding pocket, suggesting that substitutions at G67 and G98 could directly influence pocket architecture (Fig. S3A).

Among the three variants, p.G98S produced the most pronounced alteration in predicted pocket organization relative to WT, whereas p.N64S and p.G67R caused comparatively modest changes. Several canonical ATPase-pocket residues, including N38, G67, and K84, were absent from the highest-ranked G98S pocket but were recovered in a secondary predicted pocket (Fig. S3B), indicating redistribution of ATPase-associated residues rather than loss of ligand accessibility. Consistent with this, the primary G98S pocket showed markedly reduced similarity to WT (Jaccard similarity = 0.122), increasing to 0.452 when the secondary pocket was included. In contrast, G67R and N64S retained near-WT pocket organization, with Jaccard similarities of approximately 0.85. Spatial mapping of ATPase-associated residues produced a similar result. N64S largely preserved the WT arrangement, whereas G67R caused relatively localized changes, particularly around residue 67. G98S produced substantially greater displacement of several conserved ATPase residues, including N38, G67, and K84, relative to the predicted pocket center (Fig. S3C-D).

Structural superposition localized the greatest divergence between WT and G98S to residues approximately 95–100, immediately adjacent to the ATP-binding pocket (Fig. S3E-F). AlphaFold confidence in this region was relatively low (pLDDT ~40–50). Because pLDDT can reflect both prediction uncertainty and intrinsic conformational flexibility (4), we examined multiple independently generated models. Displacement of residue 98 was reproducibly observed in five G98S models, whereas corresponding N64S and G67R models remained substantially more similar to WT (Fig. S3G). This reproducibility suggests that the altered local conformation is associated with the G98S substitution rather than stochastic variation among AlphaFold predictions, while also raising the possibility that the mutation increases local conformational heterogeneity. Independent Missense3D analysis classified G98S as substitution of a buried glycine predicted to disrupt local packing and flexibility. G67R was also predicted to be structurally disruptive, primarily through severe steric clashes, but did not produce the extensive ATP-pocket reorganization observed for G98S.

Taken together, these analyses predict that p.G98S has the strongest structural effect among the variants examined. Importantly, the models do not indicate loss of ATP/$Mg^{2+}$ binding itself, but instead suggest remodeling of the ATPase environment and altered local conformational dynamics. Because ATP binding and hydrolysis drive conformational transitions required for MutL activity during mismatch repair and meiotic crossover formation, disruption of this ATPase cycle could severely compromise MLH1 function despite preserved nucleotide binding. This distinction may also have implications for the unusually severe phenotype associated with p.G98S. ATPase-defective mismatch repair proteins can retain assembly or recognition functions while failing at downstream ATP-dependent transitions. For example, the *E. coli* MutL-E32K ATPase mutant can form repair complexes but fails to couple ATP hydrolysis to downstream activation and exhibits dominant-negative activity (5). By analogy, MLH1-G98S could retain sufficient structural integrity to bind ATP and associate with PMS2 or MLH3 while failing to undergo the conformational changes necessary for productive MutLα or MutLγ function. Such incorporation of an ATPase-impaired MLH1 protein into otherwise functional complexes could potentially interfere with the activity of the WT allele and produce a dominant-negative phenotype.
